# Evaluation of neural regulation and microglial responses to brain injury in larval zebrafish exposed to perfluorooctane sulfonate

**DOI:** 10.1101/2022.09.30.510316

**Authors:** Shannon E. Paquette, Nathan R. Martin, April Rodd, Katherine E. Manz, Manuel Camarillo, Eden Allen, Hannah I. Weller, Kurt Pennell, Jessica S. Plavicki

**Author notes:** <u>Corresponding Author</u> Jessica S. Plavicki, PhD, Department of Pathology & Laboratory Medicine Brown University, 70 Ship Street, Providence, RI 02903. These authors contributed equally (Shannon E. Paquette, Nathan R. Martin). The authors have no conflicts of interest to declare.

## Abstract

**Background:** Per- and polyfluoroalkyl substances (PFAS) are biopersistent pollutants that have become global contaminants as a result of their diverse applications in commerce and industry. While some *in vitro* and epidemiological studies have explored the neurotoxic potential of perfluorooctane sulfonate (PFOS), a prevalent PFAS congener, it is unknown how developmental exposure to PFOS affects neuronal communication and other developmentally critical neural cell types, including microglia.

**Objectives:** We sought to determine the extent to which PFOS exposure disrupts brain health, neuronal activity, and neuron-microglia communication during brain development. In addition, while PFOS impairs humoral immunity, its impact on innate immune cells, including resident microglia, is unclear. As such, we aimed to determine whether microglia are cellular targets of PFOS and, if so, whether disrupted microglial development and/or function could contribute to or is influenced by PFOS-induced neural dysfunction.

**Methods:** Zebrafish were chronically exposed to either control solution (0.1% DMSO), 7 µM PFOS, 14 µM PFOS, 28 µM PFOS, or 64 µM perfluorooctanoic acid (PFOA). We used *in vivo* imaging and gene expression analysis to assess microglial populations in the developing brain and to determine shifts in microglial state. We functionally challenged microglial using a larval brain injury model and, to assess the neuronal signaling environment, performed functional neuroimaging experiments utilizing the photoconvertible calcium indicator CaMPARI. These studies were paired with optogenetic manipulations of neurons and microglia, an untargeted metabolome wide association study (MWAS), and larval swim behavior assessments.

**Results:** Developmental PFOS exposure resulted in a shift away from the homeostatic microglia state, as determined by functional and morphological changes, as well as transcriptional upregulation of the microglia activation gene *p2ry12*. PFOS-induced effects on microglia state exacerbated microglia responses to brain injury in the absence of increased cell death or inflammation. PFOS exposure also heightened neural calcium activity, and optogenetic silencing of neurons or microglia independently was sufficient to normalize microglial responses to injury. An untargeted MWAS of larval brains revealed PFOS-exposed larvae had neurochemical signatures of excitatory-inhibitory imbalance. Behaviorally, PFOS-exposed larvae also exhibited anxiety-like thigmotaxis. To test whether the neuronal and microglial phenotypes were specific to PFOS, we exposed embryos to PFOA, a known immunotoxic PFAS. PFOA did not alter thigmotaxis, neuronal activity, or microglial responses, further supporting a role for neuronal activity as a critical modifier of microglial function following PFOS exposure.

**Discussion:** Together, this study provides the first detailed account of the effects of PFOS exposure on neural cell types in the developing brain *in vivo* and adds neuronal hyperactivity as an important endpoint to assess when studying the impact of toxicant exposures on microglia function.

## INTRODUCTION

Pollution poses a global threat to environmental, ecologic, and economic health and stability. As international geologic committees consider whether we have now entered the “Anthropocene Age,” it is of increasing importance to understand the broad impacts of human-made pollutants. It is also worth emphasizing that the burden of pollution is disproportionately shared, with minority and low-income communities being more susceptible to the physical consequences of insufficient regulatory laws, industrial waste disposal, and air pollution^(1–3)^. If we are to protect our health, society, and environment, more research efforts are needed to address the costs of our past and present environmental negligence.

One major class of chemicals of increasing concern are per- and polyfluoroalkyl substances (PFAS). While PFAS were only introduced in industrial manufacturing in the mid-20^th^ century, the expansive and international utilization of PFAS has led to near universal exposure in humans^(4, 5)^. PFAS congeners are found in fire-fighting foams, commercial household products such as water-repellant fabrics, food packaging, and non-stick finishes, and are used in a variety of applications in the aerospace, aviation, and automotive industries^(6)^. Structurally, PFAS consist of fully or polyfluorinated aliphatic substances of varying carbon chain length and head groups, with longer chain length tending to be associated with increased toxicity^(7–9)^. The strength of carbon-fluorine bonds lends most PFAS to be biopersistent, bioaccumulative, and resistant to degradation^(10)^. As such, PFAS have acquired the disconcerting moniker “forever chemicals.”

Of the 4,700+ PFAS congeners^(6)^, one of the most environmentally prevalent and extensively studied is perfluorooctane sulfonate (PFOS)^(10)^. PFOS has an 8-carbon chain with a sulfonic acid head group and adversely impacts the functioning and health of several major organ systems^(11, 12)^, and is considered a metabolic, endocrine, and immune disruptor^(13–19)^. *In utero* and developmental exposure to PFOS has particularly concerning consequences on adaptive immunity. Grandjean et al. demonstrated in humans that elevated PFOS levels in infancy and early childhood significantly attenuate adaptive immune responses, curb antibody production, and limit vaccine efficacy^(20, 21)^. Such studies reinforce the necessity of understanding the impact of pollution on population health, especially considering the need to vaccinate individuals against emerging or evolving pathogens. PFOS-induced humoral immune suppression^(14, 22)^ prompted the National Toxicology Program to classify PFOS as a “presumed immune hazard to humans” in 2016^(23)^. However, the effects of PFOS on innate immunity are still largely inconclusive, with some groups finding that PFOS dampens innate cell infiltration, gene expression, or activity^(24–26)^, and others describing heightened inflammation or immune function^(27–29)^. Even less understood is the tissue-specific impact of PFOS on local immune populations, namely tissue-resident macrophages. Tissue-resident macrophages are largely yolk-sac or fetal liver-derived, heterogenous, and most notably carry out discrete, non-canonical tissue-specific functions essential for development and homeostasis^(30)^. Determining whether tissue-resident macrophages are PFAS targets, especially during development, is essential for our understanding of long-term consequences of macrophage dysregulation following PFAS exposure.

In this work, we focus on microglia, the resident immune population of the central nervous system (CNS), as a potential PFOS target. Beyond effector immune responses, microglia have a highly dynamic and diverse repertoire of homeostatic functions, including developmental pruning of extra-numerary synapses^(31, 32)^, facilitating synaptogenesis and maturation^(33)^, and regulation of neural plasticity and dendritic spine density through frequent synaptic monitoring^(34–36)^. As sentinels of the CNS, these persistent, self-renewing cells are also highly sensitive and rapidly adaptable to any changes in their environment.

Microglial dysfunction has been well documented in disease models of various neuropathological states, including Down Syndrome^(37)^, Alzheimer’s disease^(38)^, and epilepsy^(39)^. Due to the critical regulatory roles of microglia in neural health and development, there has been a concerted effort to clarify the mechanisms of bidirectional crosstalk between neurons and microglia^(35)^. However, the impact of environmental exposures on neuron-microglia interactions during developmentally sensitive periods of life, including gestation, has been largely overlooked. Some epidemiological studies suggested correlations between developmental PFOS exposure and ADHD incidence^(40, 41)^, while others found no such relationship^(42–45)^. Meanwhile in mice, a single neonatal PFOS exposure was sufficient to alter proteins required for neuronal growth and synaptogenesis, as well as cause spontaneous behavior and hyperactivity in adults^(46, 47)^. Developmental exposure to PFOS was also shown to cause hyperactive locomotor activity in larval zebrafish^(48–50)^. *In vitro* experiments using cultured neurons revealed that PFOS interacts with inositol 1,4,5-triphosphate receptors (IP_3_Rs) and ryanodine receptors (RyRs), leading to the release of intracellular calcium stores, which suggests a potential role for calcium in PFOS-induced neurotoxicity^(51)^. However, it is still not known whether the induced locomotor activity in zebrafish is attributed to neuronal hyperactivity or skeletal muscle calcium utilization^(52)^. It is also not known if potential changes in the neuronal environment following PFOS exposure can impact microglia function, and vice versa.

Herein, we utilized the versatility, transparency, and *ex vivo* development of the larval zebrafish model to determine how chronic PFOS exposure impacts microglial and neuronal function *in vivo* during development. We employed a brain injury model, functional neuroimaging, larval behavioral assays, optogenetic modulation of cell-specific membrane potential, a metabolome wide association study (MWAS), and a microglia knockout model in larval zebrafish to provide the first *in vivo* account of PFOS-induced disruption of microglial and neuronal activity. We also exposed larvae to a non-excitatory PFAS congener, perfluorooctanoic acid (PFOA), to determine whether structurally similar compounds can have distinct effects on CNS health and neural cell function.

## MATERIALS & METHODS

### Zebrafish husbandry

Zebrafish (*Danio rerio*) maintenance and experimental procedures were approved by the Brown University Institutional Animal Care and Use Committee (IACUC; 19-12-0003) adhering to the National Institute of Health’s “Guide for the Care and Use of Laboratory Animals.” Zebrafish colonies were maintained in an aquatic housing system (Aquaneering Inc., San Diego, CA) maintaining water temperature (28.5 ± 2°C), filtration, and purification, automatic pH and conductivity stabilization, and ultraviolet (UV) irradiation disinfection. Adult and larval zebrafish were sustained in a 14:10 hour light-dark cycle (Westerfield 2000). Adult zebrafish were fed once a day with GEMMA Micro (Skretting, Westbrook, ME).

Adult zebrafish (varying ages >3 months post-fertilization) were placed into 1.7 L sloped spawning tanks (Techniplast, USA) 15-18 hours prior to mating. Sexes were separated by a transparent partition. Within 2 hours of light cycle onset, the partition was removed, and zebrafish were allowed to spawn for 1 hour. Embryos were collected in fresh egg water (60 mg/L Instant Ocean Sea Salts; Aquarium Systems, Mentor, OH) and placed into 100 mm non-treated culture petri dishes (CytoOne, Cat. No. CC7672-3394). Embryonic and larval zebrafish were maintained at 28.5 ± 1°C in an incubator (Powers Scientific Inc., Pipersville, PA) up to 120 hours post-fertilization (hpf).

### Zebrafish lines

The following zebrafish lines were used in this study, either independently or in combination. To visualize green fluorescent macrophages, we used *Tg(mpeg1:EGFP)*^(53)^ zebrafish. To modulate cellular membrane potential by inducing chloride ion influx, we used the optogenetic line halorhodopsin, which is a light-gated chloride pump that is sensitive to 589 nm light^(54, 55)^. This third-generation opsin exhibits reliable membrane trafficking for uniform surface expression, is resistant to inactivation, and has step-like, potent photocurrents that are stable over long timescales^(55)^. Halorhodopsin was under the control of a Gal4-regulated upstream activating sequence (UAS) for cell-specific control, *Tg(UAS:eNpHR3.0-mCherry)*^(54, 55)^. To detect calcium activity in neurons *in vivo,* we used a transgenic line with neuron-specific expression of the calcium sensor CaMPARI (Calcium-Modulated Photoactivatable Ratiometric Integrator) under the pan-neuronal promoter *elavl3* (*Tg(elavl3:CaMPARI(W391F+V398L))^jf9^*)^(56)^. CaMPARI is a permanent photoconvertible calcium sensor that undergoes allosteric chromophore modulation from green-to-red in response to ultraviolet light, but only upon simultaneous binding of free intracellular calcium^(56)^. To achieve neuron-specific expression of a transgene of interest, we used a neuron-driven Gal4 line, *Tg(elavl3:Gal4-VP16)*^(57)^. Likewise, to achieve macrophage-specific expression of a transgene, we used a macrophage-driven Gal4, *Tg(mpeg1:Gal4FF)^gl25^*^(53)^. To visualize neurons and macrophages simultaneously, we used a photoconvertible pan-neuronal transgenic line in which the neurons express a photoconvertible protein, Kaede, that converts from green-to-red in response to blue light, *Tg(HuC:Kaede)*^(58)^. To visualize macrophages in the red channel as a control experiment for the optogenetic stimulation study, we used a transgenic line with UAS-driven nitroreductase fused to mCherry, *Tg(UAS:nfsB-mCherry)*^(59)^ and crossed these fish with the macrophage-Gal4 line. To determine whether or not PFOS-induced microglial dysfunction during early larval development influenced neuronal excitation and behavior, we used the embryonic macrophage mutant line targeting the gene *irf8* (TALE-NT 2.0; frameshift mutation of the st96 allele)^(60)^. Irf8 mutants lack all embryonic-derived macrophage populations, including microglia^(60)^. This line was kindly provided to us by Dr. William S. Talbot of Stanford University.

### Perfluorooctanesulfoinc acid (PFOS) and perfluorooctanoic acid (PFOA) exposure

Perfluorooctane sulfonic acid (PFOS; Sigma-Aldrich, Cat. No. A-5040) and perfluorooctanoic acid (PFOA; Sigma-Aldrich, Cat. No. 171468) were prepared by dissolving the powdered compounds in 100% DMSO. PFOS and PFOA stock concentrations were verified using LC-HRMS (certified reference standards and internal standards were purchased from Wellington, Ontario, Canada). Upon verification of the stock concentration, we found that the PFOS solution contained the following impurities: 4409.49 ng/L of PFHpS, 4588.56 ng/L of PFHxS, and 7493.40 ng/L of PFNS. PFOS and PFOA stock solutions were diluted by a factor of 5000x in a mixture 1:1 methanol:water and 2 mM ammonium acetate to accommodate the detection range of the LC-HRMS. LC/MS grade water and methanol were purchased from Honeywell (Muskegon, MI 49442). Ammonium acetate solution (5 M) was purchased from Millipore Sigma (Burlington, MA 01803).

Timed spawns of relevant transgenic zebrafish crosses were performed for 1 hr. Embryos were collected and screened for embryo quality at 4 hpf. Healthy embryos were placed in 24-well plates at a density of 3 embryos per well. Prior to treatment, PFAS compounds were diluted in egg water to the final concentration of interest. Egg water containing 0.1% DMSO was used as vehicle control. Embryos were dosed with 2 mL of diluted PFOS solution or vehicle control per well at 4 hpf. The 24-well plates were sealed with parafilm to limit evaporative loss and placed in an incubator (28.5 ± 1°C). Embryos were dechorionated at 24 hpf and statically exposed until the experimental timepoint of interest. PFOS body burden was assessed at 48 hpf and 72 hpf as described in the section “Targeted Analysis of PFOS Using Liquid Chromatography High Resolution Mass Spectrometry” below.

We selected a range of PFOS concentrations based on previously published developmental toxicology research that provided important information regarding mortality and gross morphological changes in response to exposure in larval zebrafish at similar timepoints^(61, 62)^. The survival rate following exposure to the three highest concentrations of PFOS used in the study (14 µM PFOS, 28 µM PFOS, and 56 µM PFOS) was determined between 24 hpf (1 dpf) and 120 hpf (5 dpf). At 120 hpf, there was no significant difference in larval survival in the 14 µM PFOS-exposed group. However, significant reductions in survival were first detected at 96 hpf in the 28 µM and 56 µM PFOS-exposed groups (Table S1). As such, larvae exposed to 7 µM and 14 µM PFOS were assessed until 5 dpf, while larvae exposed to 28 µM or 56 µM PFOS were assessed until 3 dpf.

### Brain injury model

At 3 dpf, larval zebrafish were anesthetized in 0.02% tricaine-s solution (Syndel, Ferndale, WA) and restrained dorsal-side-up in 2% agarose (Fisher Scientific, Cat. No. BP160-100). Larvae were injured anteriorly at the right telencephalon with a 9 um OD pulled glass capillary needle. This method was an adaptation for larval fish^(63)^. For time-lapse imaging, larvae were immediately mounted dorsally on a 35 mm glass bottom microwell dish (MatTek, Part No. P35G-1.5-14-C) in 2% low-melting agarose (Fisher Scientific, Cat. No. BP160-100) surrounded by egg water. Multi-slice projection images of the forebrain and optic tectum at various timepoints post-injury, as well as time-lapse videos composed of 10-minute imaging intervals, were captured using a Zeiss LSM 880 confocal microscope at 20x magnification. Area of microglia response was measured using Zen Blue (Zeiss).

### Microglial cell quantification

Adult transgenic zebrafish expressing *Tg(HuC:Kaede)* and *Tg(mpeg1:EGFP)* were crossed to generate *Tg(HuC:Kaede; mpeg1:EGFP)* fish, which had pan-neuronal expression of the photoconvertible protein kaede as well as green macrophages. Embryos were exposed to PFOS as described. At 3 dpf, larvae were fixed in 4% paraformaldehyde (PFA, Sigma-Aldrich, Cat. No. P6148) for 18-24 hours at 4°C. Post-fixation, larvae were washed 3 times in PBS-T (phosphate buffered solution + 0.6% Triton-X 100) (Sigma-Aldrich, Cat. No. X100). PBS-T was removed and VECTASHIELD Mounting Media (Vector Labs, Cat. No. H-1000) was added to the samples. Samples incubated for 5 minutes at room temperature, then were gently mixed and placed in -20°C until imaging. The larvae were removed from VECTASHIELD and mounted in 2% low-melting agarose (Fisher Scientific, Cat. No. BP160-100) in 35mm glass bottom microwell dishes (MatTek, Part No. P35G-1.5-14-C). Prior to image acquisition, the kaede fluorophore was photoconverted from green to red fluorescence using 405 nm light (about 1-minute exposure) on the Zeiss LSM 880 confocal microscope. Microglia were defined as macrophages (mpeg1+ cells) amongst and in contact with differentiated neurons (HuC+ cells) in the forebrain, optic tectum, and hindbrain. The entirety of the zebrafish brain was imaged at 20x magnification (2 tile panels with 10% overlap). Microglia were counted by panning through confocal z-stacks using FIJI/ImageJ. Immune cells in contact with HuC+ differentiated neurons were counted as parenchymal cells (i.e. microglia). Surface cells in contact with differentiated neurons that failed to extend into the parenchyma were defined as non-parenchymal cells (i.e. macrophage). Location and proximity of mpeg1+ cells were confirmed using z-stack orthogonal views.

### Microglial morphology quantification following PFOS exposure

Microglial morphology was performed on high-resolution confocal micrographs of the optic tectum in fixed transgenic zebrafish larvae expressing either *Tg(mpeg1:EGFP), Tg(mpeg1:Gal4FF;UAS:eNpHR3.0-mCherry),* or *Tg(mpeg1:GalFF;UAS:nfsb-mCherry).* Individual microglia were isolated based on their specific z-stack range, to prevent analysis of overlapping cells, using FIJI/ImageJ. Z-slices were also processed to rescale intensity (compensating for loss of intensity in deeper tissue) and modestly smoothened using Gaussian blur with a sigma radius of 1 to better define cellular extensions and reduce signal noise. Each cell was individually outlined, and morphological parameters measured with CellProfiler software (version 3.1.9) or FIJI/ImageJ. Area, perimeter, and cell shape (perimeter-to-area ratio) were quantified.

### Quantitative RT-PCR

Larval zebrafish heads were collected at 3 dpf. To collect, larvae were placed on ice for 4 minutes to immobilize swim activity but prevent pain or distress during collection^(64)^. The heads were removed by cutting at the base of the hindbrain at a 45° angle, limiting collection of the heart and pericardium. Heads were pooled (n = 10) and flash frozen in liquid nitrogen. Four independent experimental replicates were collected per treatment group. RNA isolation and purification was carried out using the Rneasy Plus Kit (QIAGEN). cDNA synthesis was achieved using the SuperScript IV Reverse Transcriptase First-Strand Synthesis System kit (Invitrogen, Cat. No. 18091050). qRT-PCR for genes of interest was performed with 7.5 ng/uL of cDNA using the ViiA7 Real Time PCR System (Applied Biosystems). Gene targets were detected by using either pre-designed TaqMan probes (Thermo Fisher Scientific) or custom primers with Power Track SYBR Green Master Mix (Thermo Fisher, Cat. No. A46012). See list of probes and primers for qRT-PCR and genotyping in Table S2.

### Genotyping *irf8* mutants

*irf8*^st^^96/st96^ were generated by Shiau et al. via TALEN-targeting^(60)^. After swim behavior testing or CaMPARI imaging, whole larvae were placed directly in 50 uL of DNA lysis buffer (10 mM Tris pH 8.4 (Fisher, Cat. No. BP152-500), 50 mM KCl (Sigma, Cat. No. P9333), 1.5 mM MgCl2 (Sigma, Cat. No. M0250), 0.30% Tween-20 (Fisher, Cat. No. BP337), 0.30% Igepal/NP-40 (Sigma, Cat. No. 56741)) to be used for genotyping. Larvae were incubated in DNA lysis buffer for 20 minutes at 94°C, cooled to 55°C, then 5 uL of 10 mg/mL of proteinase K (Millipore Sigma, Cat. No. 70663-4) was added. All temperature specific reactions were carried out in the ProFlex PCR System (Applied Biosystems). Samples were then incubated for 1 hour at 55°C, followed by 20 minutes at 94°C, and held at 4°C until the PCR reaction. The PCR reaction was performed with 2 uL of DNA and 18 uL of master mix (diH2O, 10x Taq Buffer (Thermo Scientific, Cat. No. EP0404), 2.5 mM dNTP (Thermo Scientific, Cat. No. R1121), 25 mM MgCl2 (Thermo Scientific, Cat. No. AB0359), 10 uM forward primer (IDT), 10 uM reverse primer (IDT), DMSO (Millipore Sigma, Cat. No. D8418), and Taq DNA polymerase (Thermo Scientific, Cat. No. EP0404)). The PCR reaction conditions were as follows: 1 minute at 95°C; 35 cycles of 30 seconds at 95°C, 45 seconds at 68°C, and 30 seconds at 72°C; then 7 minutes at 72°C; and hold at 4°C. The *irf8* exon 1 fragment was amplified using primers listed in Table S2. As this *irf8* mutant has a frameshift mutation at an AvaI site, the PCR product was used for restriction digest with AvaI (New England BioLabs, Cat. No. R0152L) to identify the presence of the mutation. For the restriction digest reaction, 10 uL of PCR product and 10 uL of digest master mix (diH2O, rCutSmart Buffer, and AvaI) were combined and run on the thermocycler for 20 minutes at 37°C and cooled to 4°C. The digest reaction product was run in a 2% agarose gel containing RedSafe nucleic acid staining solution (Bulldog Bio Inc, Cat. No. 21141). The gel results were visualized using the Bio-Rad Gel Doc XR+.

### Acridine orange staining

Cell death was determined using the vital dye acridine orange (AO) in live zebrafish embryos. AO is a cell permeable stain that selectively intercalates uncoiled nucleic acid present in apoptotic cells^(65)^. Zebrafish embryos were injured at 3 dpf and placed in 5 ug/mL acridine orange (ThermoFisher, Cat. No. A1301) diluted in egg water for 20 minutes. Zebrafish were then washed in egg water for 15 minutes, refreshing the solution every 5 minutes. Live zebrafish were mounted in 2% agarose and immediately imaged on a Zeiss LSM 880 confocal microscope. For 1 hour-post injury (hpi) analysis, staining took place from 40 minutes to 1 hpi. For 4 hpi analysis, staining took place from 3 hours and 40 minutes to 4 hpi.

### CaMPARI photoconversion and image acquisition

*Tg[elavl3:CaMPARI(W391F+V398L)]^jf9^* embryos were screened and the brightest (highest expressing) were selected for toxicant exposure and subsequent photoconversion. At either 3 dpf or 5 dpf, CaMPARI-positive larvae were individually placed into a modified 1-well dish (15 mm diameter) containing PFOS or 0.1% DMSO. The dish was placed onto a constructed pedestal within a DanioVision Observation Chamber (Noldus, Wageningen, The Netherlands) adapted with optogenetics components (Prizmatix, Southland, MI). Use of the pedestal decreased working distance from the LED light source to the free-swimming larva, concordantly increasing light intensity. Cells expressing CaMPARI with high calcium content will photoconvert (green-to-red) in the presence of 405 nm light. Photoconversion was performed by exposing zebrafish to 405 nm (135 mW/cm^2^) wavelength light for 1 minute. Live zebrafish larvae were then anesthetized in 0.02% tricaine-s (MS-222) and mounted in 2% low-melting agarose in 35mm glass bottom microwell dishes. Confocal z-stacks were acquired on a Zeiss LSM 880 confocal microscope and maximum intensity projections generated in Zen Black (Zeiss) representing a global snapshot of neural activity. Image parameters were set during acquisition of the first image and maintained for the duration of each experiment.

### CaMPARI analyses (R/G ratio calculations)

CaMPARI photoconversion from green-to-red during exposure to 405 nm wavelength light was used to determine neuronal calcium levels following PFOS exposure at 3 dpf or 5 dpf. To assess the level of photoconversion, maximum intensity projections were imported into FIJI/ImageJ. Brain regions of interest including forebrain (FB), habenula (H), optic tectum (OT), cerebellum (Ce), hindbrain (HB), and whole brain (WB) were manually selected and integrated density of red and green was measured. Measurements were blank corrected by selecting 3 separate background regions of each image (Equation 1). The ratios of corrected integrated density were then used to determine R/G ratio for each respective image (Equation 2). R/G ratios were then normalized to the average control R/G ratio within a particular experiment (Equation 3).

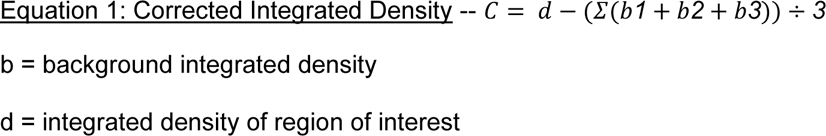

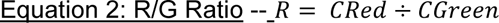

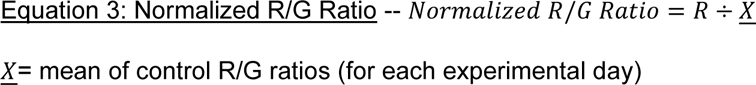

### Pentylenetetrazol (PTZ) exposure

For validation of CaMPARI functionality, 3 dpf *Tg(elavl3:CaMPARI(W391F+V398L))^jf9^* larvae were exposed to 10 mM pentylenetetrazol (PTZ) for 10 minutes. PTZ was washed out 3 times with egg water immediately followed by CaMPARI photoconversion, as described. For assessment of microglia response to injury following PTZ exposure, larvae were exposed to control solution, 28 µM PFOS, or 5 mM PTZ at 3 dpf. Larvae were injured in the right telencephalon, as described, and placed back in their original dosing solutions for 4 hours. At 4 hpi, larvae were mounted in 2% agarose in a 35mm glass bottom imaging dish and multi-slice projection images of the forebrain and optic tectum were captured using a Zeiss LSM 880 confocal microscope at 20x magnification. Area of microglia response was measured using Zen Blue (Zeiss).

### Light/dark behavioral assay

Behavioral assessments were performed in 24-well plates (ThermoFisher, Cat. No. 144530) using a DanioVision Observation Chamber with EthoVision XT live-tracking software (Noldus, Wageningen, The Netherlands). Larval zebrafish were exposed to either control solution, 7 µM, 14 µM, or 28 µM PFOS at 4 hpf, as described above. Approximately 18-24 hours prior to behavioral assessment, larvae were transferred into individual wells of the same dosing solution. All dosing solutions were prepared and plated on the same day as the initial larval exposure to maintain “chronic” exposure conditions, rather than introducing the larvae to a fresh dosing solution. All light/dark behavioral assays were performed between 8 AM and 12 PM to prevent behavioral changes attributable to circadian differences. Briefly, zebrafish were placed into the DanioVision observation chamber and acclimated during a 15-minute dark cycle, followed by a 5-minute light cycle (Light 1), a 5-minute dark cycle (Dark 1), another 5-minute light cycle (Light 2), and completed with a 15-minute dark cycle (Dark 2). Behavioral experiments lasted a total of 45 minutes.

Total distance moved (mm) during light/dark cycles was quantified using the EthoVision XT software as a measure for hyperactivity. Anxiety-like behavior following PFOS exposure was assessed by monitoring larval well location (center vs. edge) over the course of the assay. Defined center and edge regions each constituted 50% of total well area. Edge versus center preference was quantified using EthoVision XT. Center avoidance data was normalized relative to controls for each experimental replicate.

### Optogenetic manipulation in Noldus behavioral unit

To stimulate the optogenetic channel halorhodopsin in the transgenic lines *Tg(elavl3:Gal4;UAS:eNpHR3.0-mCherry)* and *Tg(mpeg1:Gal4ff;UAS:eNpHR3.0-mCherry),* screened larvae were placed a DanioVision observation chamber (Noldus, Wageningen, The Netherlands) outfitted with a 570 nm wavelength laser (Prizmatix, Southland, MI). Halorhodopsin’s peak excitation wavelength is 589 nm, though because its excitation spectrum is broad, halorhodopsin is also responsive to 570 nm. Halorhodopsin^+^ neurons or microglia were stimulated with 570 nm light for 4 hours then immediately imaged for cell morphology or injury response.

### Targeted analysis of PFOS using liquid chromatography high resolution mass spectrometry (LC-HRMS)

Body burden was assessed at 48 hpf and 72 hpf to determine the level of uptake of PFOS in the larvae, as well as to determine the extent to which the detected PFAS impurities (PFNS, PFHpS, PFHxS) were present in the larvae. Body burden results can be found in Table S3.

#### Chemicals

Certified PFOS, PFNS, PFHpS, and PFHxS and their respective isotope labelled reference standards were purchased from Wellington Laboratories (Ontario, Canada). LC/MS grade water and methanol were purchased from Honeywell (Muskegon, MI 49442). Ammonium acetate solution (5 M) was purchased from Millipore Sigma (Burlington, MA 01803).

#### Sample Extraction

Three technical replicates of ten larvae were pooled, flash frozen in liquid nitrogen, and stored in a -80°C freezer until extracted and defrosted at 20°C. 1 mL methanol was added to the centrifuge tube containing the embryos. The samples were sonicated for 90 minutes, vortex mixed 1 minute, and allowed to reach equilibrium for 3 hours at 20°C. Samples were then centrifuged at 3,000 rpm for 10 minutes. The following was added to an LC analysis vial: 50 µL of the methanol extract, 10 µL labelled PFOS internal standard, and 440 µL of a mixture containing 1:1 methanol:water and 2 mM ammonium acetate.

Targeted analysis of PFOS was performed using a Thermo Liquid Chromatography (LC) Orbitrap Q Exactive HF-X mass spectrometer (MS) equipped with a Thermo Vanquish UHPLC system. Mobile phase A contained 2 mM ammonium acetate in water and mobile phase B contained 2 mM ammonium acetate in methanol. 20 µl of each sample extract was injected in triplicate and separated in a Thermo Hypersil Gold Vanquish C18 column (50 mm X 2.1 mm x 1.9 µm) at a constant temperature of 60°C. PFOS was eluted from the column at a constant flow rate of 0.4 mL/min using a mobile phase gradient as follows: equilibration with 10% B for 1 minute, followed by a gradient ramp from 10% B to 95% B over 4 minutes and held for 2 minutes, and back to 10 % B over 1 minute and held for 2 minutes (total run time 9 minutes, data collected from 0.6 to 8.5 minutes. The MS was operated in full scan dd-MS^2^ mode (70 NCE) with an inclusion list for PFOS. Ionization was performed in negative mode with an ionization window of 1.0 m/z, sheath gas flow rate of 40, auxiliary gas flow rate of 10, sweep gas flow rate of 2, spray voltage of 2.7 kV, 310°C capillary temperature, funnel RF level of 35, and 320°C auxiliary gas heater temperature. Ions were further fragmented in the HCD collision cell filled with N2 (produced by a Peak Scientific Nitrogen Generator, Genius NM32LA). For the full-scan, the Orbitrap was operated with a resolution of 120,000, automatic gain control (AGC) of 3e6, and maximum dwell time of 100ms. For dd-MS2, the Orbitrap was operated with a resolution of 15,000, AGC of 2e5, and maximum dwell time of 400ms.

Four ions were monitored for PFOS, including 498.9302 m/z (quantifying), 79.9573 m/z (confirming), 98.9556 m/z (confirming), and 82.9607 m/z (confirming). The retention time of PFOS was 5.18 min. Quantification was performed in TraceFinder 5.0 General with external seven-point calibration curve prepared by serial dilutions of the calibration standard. Limits of detection (LOD) were determined by injecting a calibration 7 standard seven times and using Equation 4:

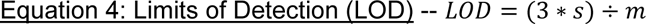

where s is the sample standard deviation and m is the calibration curve slope. The resulting PFOS LOD was 33.87 ppt.

### Untargeted metabolome wide association study (MWAS)

#### High resolution metabolomics

The sample extracts were re-analyzed using LC-HRMS to collect untargeted metabolomics data. A 10 µL volume was injected in triplicate onto on the Thermo LC-Orbitrap system described above. Two chromatography separation methods were used, normal and reverse-phase. The normal-phase LC was performed with a HILIC column (Thermo Syncronis HILIC 50 mm X 2.1 mm x 3 µm) at a constant temperature of 25°C. Mobile phase A contained 2 mM ammonium acetate in acetonitrile and mobile phase B contained 2 mM aqueous ammonium acetate. Metabolites were eluted from the column at a constant flow rate of 0.2 mL/minute using a solvent gradient as follows: equilibrate with 10% B for 1 minute, increase to 65% B for 9 minutes and hold for 3 minutes, decrease to 10% over 1 minute and hold for 1 minute. The reverse-phase LC was performed with a C18 column (Thermo Hypersil Gold Vanquish, 50 mm X 2.1 mm x 1.9 µm) at a constant temperature of 60°C. Mobile phase A contained 2 mM aqueous ammonium acetate and mobile phase B contained 2 mM ammonium acetate in acetonitrile. Metabolites were eluted from the column at a constant flow rate of 0.5 mL/minute using a mobile phase gradient as follows: equilibration with 2.5% B for 1 minute, increase to 100% B over 11 minutes and held for 2 minutes, and back to 2.5% B over 1 minute and held for 1.5 minutes (total run time 16.5 minutes, data were collected from 0.05 to 12.5 minutes). For both normal and reverse-phase LC, the MS was operated in full scan mode with 120,000 resolution, automatic gain control of 3 x 10^6^, and maximum dwell time of 100 ms. Electrospray ionization was conducted in positive mode for normal-phase and negative mode for reverse phase LC. Ionization was performed at a sheath gas flow of 40 units, auxiliary gas flow of 10 units, sweep gas flow of 2 units, spray voltage of 3.5 kV, 310°C capillary temperature, funnel radio frequency (RF) level of 35, and 320°C auxiliary gas heater temperature.

#### Metabolomics data analysis

Data files were converted from *.raw files to *.cdf files using Xcalibur file Converter, and then processed in R packages apLCMS^(66)^ and xMSanalyzer^(67)^ to produce m/z feature tables. Feature intensities were normalized by log2 transformation. Association of metabolite feature with PFOS exposure was assessed using a t-test (*P* < 0.05) in comparison to the control samples. The *P*-values were adjusted using Benjamini and Hochberg with a false discovery rate (FDR) ≤ 20% to control for Type I errors in multiple comparisons. Significant metabolites were analyzed for pathway enrichment using MetaboAnalystR^(68)^ using the zebrafish mummichog curated model, which includes the KEGG, BiGG, and Edinburgh maps. All metabolomics data analysis was performed in R (version 4.0.2).

### Statistical analyses and reproducibility

Each experiment was carried out in at least three independent experimental replicates. An experimental replicate was considered a cohort of zebrafish that were spawned on separate days and, when applicable, dosed with separate freshly prepared dosing solutions. Dosing groups for each experimental replicate were composed of siblings, such that sibling controls could be compared to dosed siblings. For each graph, datasets were tested for normality using a Shapiro-Wilk test and by plotting a QQ plot. If the data were normally distributed, parametric tests were performed, when applicable. If the data were not normally distributed, non-parametric tests were performed, when applicable. When comparing two independent groups, unpaired parametric t-tests with Welch’s correction were performed if the data were normally distributed, or Mann Whitney U tests (non-parametric) were performed if the data was not normally distributed. In cases where there were 3 or more groups with one independent variable and the data were normally distributed, a Welch ANOVA with post-hoc Dunnett’s T3 multiple comparisons test was performed. In cases where there were 3 or more groups with one independent variable and the data were not normally distributed, a Kruskal-Wallis test (non-parametric ANOVA) with Dunn’s test for multiple comparisons was performed. In cases where there were 3 or more groups with two independent variables and the mean of each group was compared to the mean of every other group, a two-way ANOVA with a post-hoc Tukey multiple comparisons test was performed. In cases where there were 3 or more groups with two independent variables and the mean of two selected groups were selected for comparison, a two-way ANOVA with a post-hoc Sidak multiple comparisons test was performed. All statistical analyses were performed using GraphPad Prism 9 (Dotmatics, San Diego, CA).

### Data availability

All data is available in the main text and supplementary materials. All targeted and untargeted metabolomics files will be deposited into Metabolomics Workbench.

## RESULTS

### Microglial morphology and response to brain injury following chronic PFOS exposure

Whether PFOS affects microglia development and function *in vivo* is not yet known. Given the critical developmental and modulatory roles of microglia in the CNS, we sought to determine whether embryonic exposure to PFOS had a direct effect on microglia colonization of the developing brain. Transgenic zebrafish embryos with macrophage-specific GFP expression under the *mpeg1* promoter (*Tg(mpeg1:EGFP)*)^(53)^ were chronically exposed to either a control solution (0.1% DMSO) or 28 μM PFOS from 4 hours post-fertilization (hpf) until 3 days post-fertilization (dpf) (Figure 1A). Using liquid chromatography with high-resolution mass spectrometry (LC-HRMS), we validated our exposure solutions and determined that 28 μM PFOS resulted in a total body burden of 70.46 ± 2.72 ng/embryo at 3 dpf (Table S3). Next, we examined microglial number and morphology at 3 dpf, a time point at which no significant differences in mortality were observed across the tested concentration range (Table S1). Non-parenchymal, parenchymal, and total microglia number was not affected at 3 dpf between the control and 28 μM PFOS-exposed larvae (Figure S1A-C). While microglia number was unchanged, cell morphology was significantly different. PFOS-exposed microglia acquired a less ramified and more amoeboid cell shape (Figure 1D-E” vs 1I-J”). This corresponded to a significant reduction in cell area (Figure 1N), cell perimeter (Figure 1O) and the acquisition of a more circular cell shape as calculated by a decrease in the perimeter-to-area ratio (Figure 1P). Relative mRNA analysis for the gene *p2ry12*, a G-protein coupled receptor for ATP directly involved in microglia activation and migration behavior^(69–71)^, was also significantly upregulated in the brains of PFOS-exposed larvae (Figure 1Q), further supporting that PFOS exposure was affecting microglial state. To determine whether the phenotypic and transcriptional changes were accompanied by functional changes, we tested whether pollutant-induced microglia activation affected the ability of microglia to respond to minor brain injury using an established zebrafish injury model^(72, 73)^. At 3 dpf, larvae were injured at the right hemisphere of the telencephalon using a pulled glass needle (OD 9 um; Figure 1B) and microglia recruitment to the injury site was monitored for the first 4.5 hours post-injury (hpi; Figure 1C). PFOS-exposed larvae had significantly more microglia recruit to the injury site in the first 4.5 hpi compared to sibling controls (Figure 1R; Supplemental videos 1 versus 2). Additionally, the area of microglial response in exposed larvae was significantly expanded at 4 hpi (Figure 1G vs 1L; Figure 1S), which persisted up to 12 hpi (Figure 1H vs 1M; Figure 1T). A major function of microglia following brain injury is the removal of damaged cells and cellular debris to prevent excessive inflammation, which is deleterious to brain health^(74)^. Using the nucleic acid stain acridine orange in live larvae, we asked whether an increased incidence of cell death was driving the microglia response to injury in pollutant-exposed larvae; however, PFOS-exposure did not result in more cell death in the whole brain nor at the injury site at 1 or 4 hpi (Figure S1D-H). Moreover, relative mRNA expression of the proinflammatory cytokines *il1β*, *tnfα*, and *il6* were unchanged in larval brains following exposure (Figure S1I).

**Figure 1.**
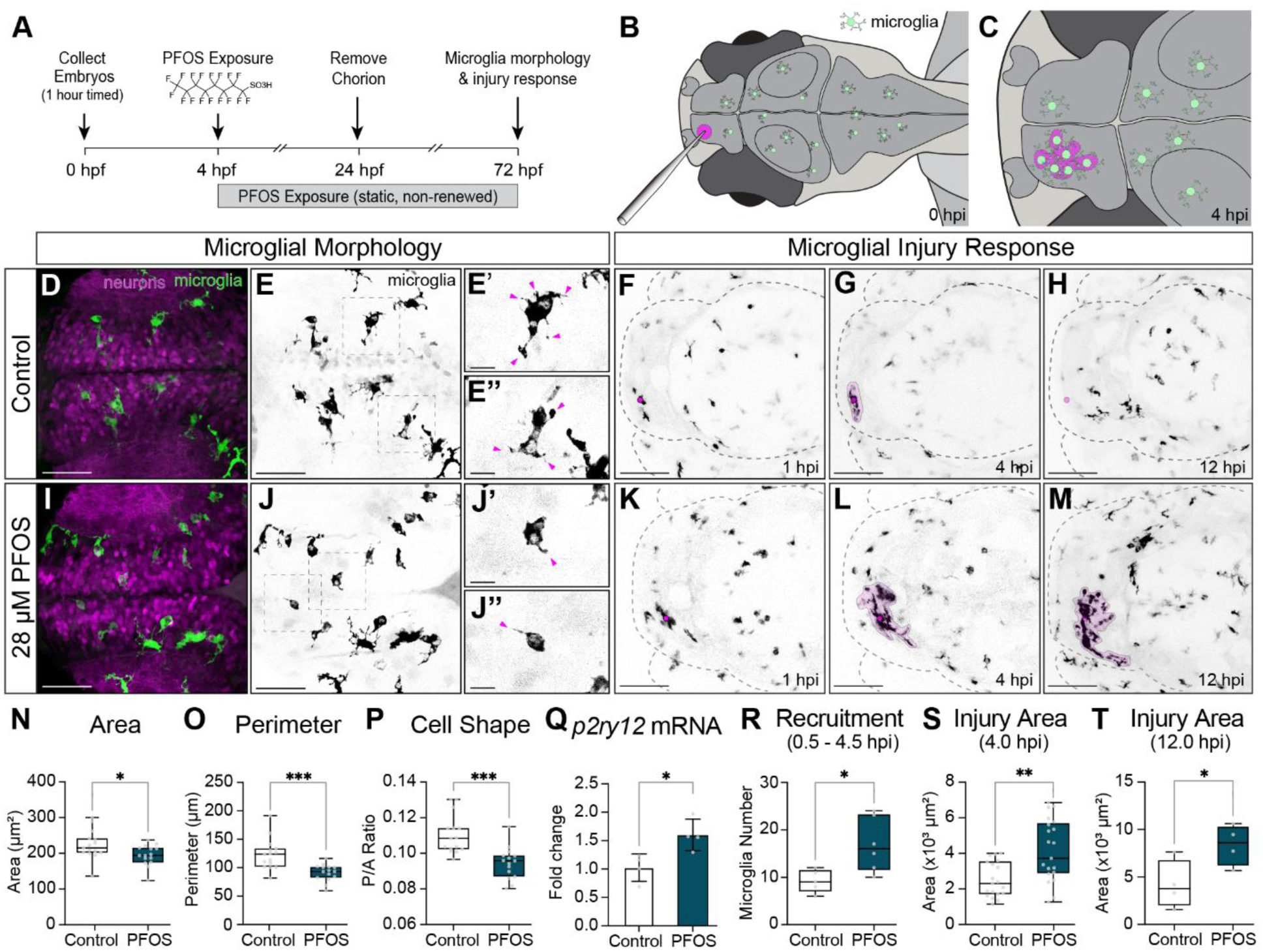
Microglia morphology and response to minor brain injury in 3 dpf larvae exposed to 28 µM PFOS. (A) Exposure paradigm: zebrafish embryos from *Tg(mpeg1:EGFP)* adults were collected after a 1- hour timed spawn. At 4 hpf, embryos were dosed with 0.1% DMSO (Control) or 28 µM PFOS. Treatment solutions were static and not renewed. Embryos were dechorionated at 24 hpf, and imaging and injury experiments were conducted at 3 dpf. (B) Schematic of larval brain injury: a pulled glass needle (OD 9 um) was used to puncture the right telencephalon of larval zebrafish. (C) Hypothetical schematic of microglia response 4 hours post-injury (hpi). (D-E”) Homeostatic, non-activated microglia of 3 dpf control larvae have several projections emanating from their cell bodies. (F-H) Control microglia respond to brain injury, with visible accumulation at the injury site at 4 hpi that mostly clears by 12 hpi. (I-J”) Microglia of 3 dpf larvae exposed to 28 µM PFOS become phenotypically activated, with rounder cell bodies and reduced projections. (K-M) In PFOS-exposed larvae, microglia robustly respond to brain injury by 4 hpi and are still maintained at the site at 12 hpi. Quantifications of the images reveal that PFOS-exposed microglia (N) area (*P* = 0.0342) and (O) cell perimeter (*P* = 0.0006) are significantly reduced. (P) The perimeter-to-area ratio (*P* = 0.0002) is also significantly decreased, indicative of a rounder cell morphology. n = 15 fish per group (3-18 cells counted per fish). (Q) qRT-PCR for the microglia activation gene, *p2ry12*, was significantly upregulated in isolated heads of PFOS-exposed larvae (*P* = 0.0211; n = 10 pooled heads per sample). PFOS exposure resulted in a significant increase in microglia recruitment in the first 4.5 hpi (*P* = 0.0208; n = 5-6 per group), as well as increased response area at (S) 4 hpi (*P* = 0.0037; n = 19-20 per group) and (T) 12 hpi (*P* = 0.0467; n = 4 per group). Confocal micrographs at 40x magnification (D-E” & I-J”) or 20x magnification (F-H; K-M). Box plot limits represent 25^th^ to 27^th^ percentile, with the midline representing the median. See Table S5 for additional statistical details.

### Microglia morphology and response to brain injury following optogenetic reversal of microglia activation state in PFOS-exposed larvae

To determine the extent to which microglial state contributed to the heightened microglial response to injury, we next asked whether PFOS-induced changes in microglia state could be reversed *in vivo*, and if this reversal of state was sufficient to normalize the injury response. Since the homeostatic microglia membrane potential is largely maintained by chloride channel currents^(75)^, we utilized transgenic zebrafish expressing the light-gated chloride pump halorhodopsin (eNpHR3.0) specifically in microglia (*Tg(mpeg1:Gal4FF;UAS:eNpHR3.0-mCherry)*) (Figure 2A). Larvae with halorhodopsin+ microglia were exposed to 28 μM PFOS, injured at 3 dpf, and exposed to 570 nm light for 4 hours (Figure 2B,C) to induce the halorhodopsin construct specifically in mpeg+ cells. We confirmed the PFOS-induced changes in microglial morphology in unstimulated PFOS-exposed larvae (Figure 2D-E”) and compared the microglia morphology of these unstimulated larvae to PFOS-exposed larvae stimulated with 570 nm light. Halorhodopsin stimulation of PFOS-exposed microglia produced a significantly more ramified morphology (Figure 2H-I”). Additionally, while stimulation had no effect on microglia area (Figure 2L), it significantly increased the cell perimeter (Figure 2M) and perimeter-to-area ratio (Figure 2N), supporting the transition from an amoeboid to a ramified state. We also confirmed 570 nm stimulation alone had no impact on microglia morphology by stimulating and assessing microglia from PFOS-exposed larvae lacking halorhodopsin expression, *Tg(mpeg1:Gal4FF;UAS:nfsb-mCherry)* (Figure S2). We next tested whether optogenetic modulation of microglia state was sufficient to rescue their responses to brain injury. Indeed, while unstimulated larvae had exacerbated microglial responses to injury (Figure 2F vs 2G), stimulation of halorhodopsin in PFOS-exposed microglia significantly reduced the microglia response (Figure 2G vs 2K; area of response quantified in Figure 2O). These data suggest that modulation of microglia activity state alone is sufficient to normalize their functions following toxicant exposure.

**Figure 2.**
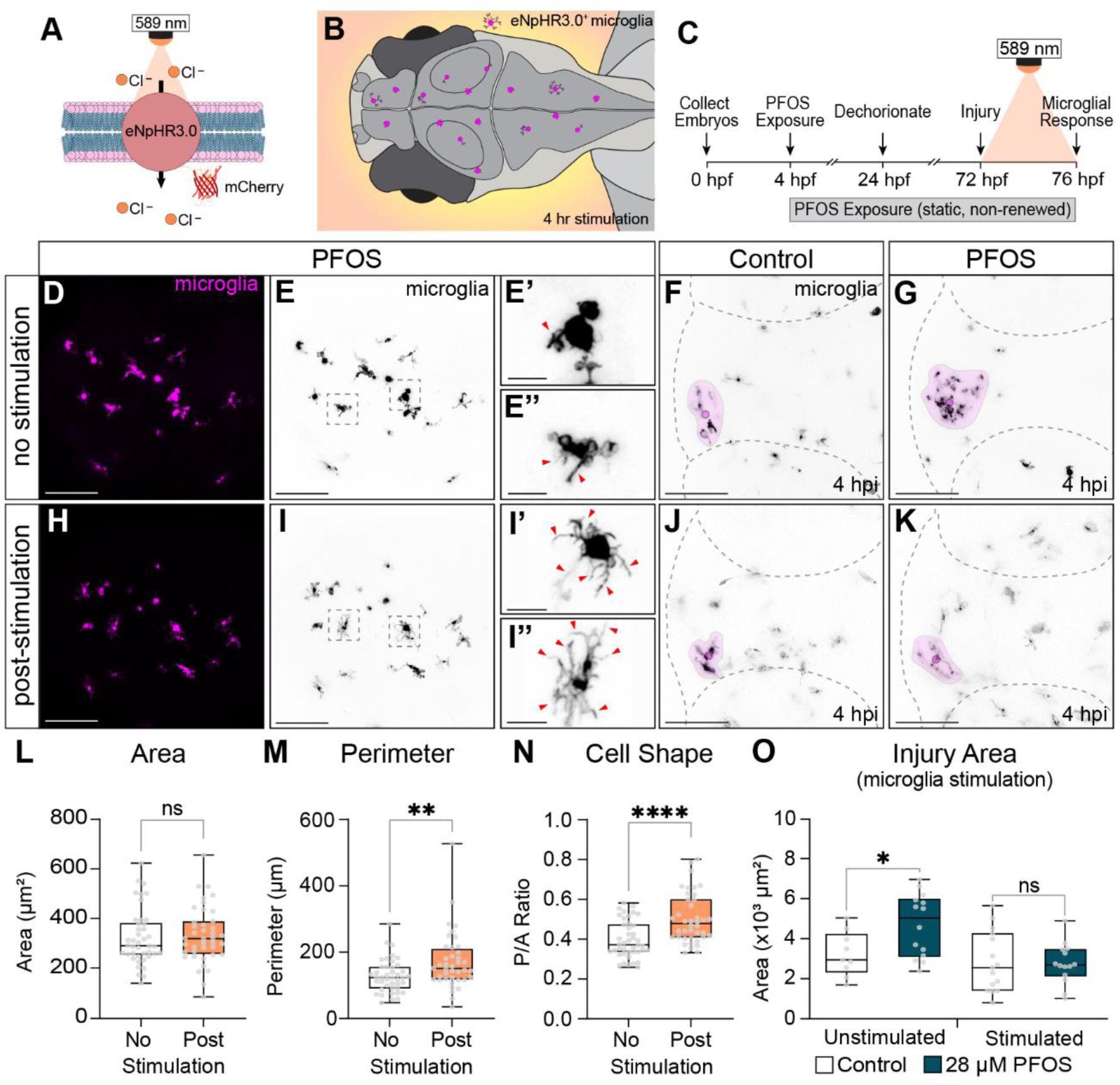
Optogenetic modulation of PFOS-exposed microglia. (A) Schematic of halorhodopsin: Optogenetic modulation of microglia electrical state is achieved via photo-stimulation of the light-gated chloride pump, halorhodopsin (eNpHR3.0). eNpHR3.0 is most responsive to 589 nm wavelength light. (B) eNpHR3.0 was driven under a pan-macrophage promoter (*Tg(mpeg1:GalFF;UAS:eNpHR3.0-mCherry)*) to achieve optogenetic control of microglia in zebrafish larvae. (C) Experimental paradigm: At 72 hpf, injured or uninjured zebrafish were stimulated for 4 hours with 589 nm light in the enclosed Noldus DanioVision Behavior Unit. (D-E”) Unstimulated halorhodopsin^+^ microglia of 3 dpf 28 µM PFOS-exposed larvae were rounded with few projections. As shown in Figure 1, (F) unstimulated control microglia were responsive to minor injury at 4 hpi, though (G) unstimulated PFOS-exposed microglia had a significantly heightened response. (H-I”) Following 4-hour stimulation of halorhodopsin^+^ in PFOS-exposed larvae, microglia became more ramified. (J, K) Stimulation of halorhodopsin^+^ microglia following PFOS exposure also normalized the microglia response to injury. (L) Stimulated microglia area was unchanged, but (M) they had significantly increased cell perimeter (*P* = 0.007) and (N) an increased perimeter-to-area ratio (*P* < 0.0001), indicative of a more ramified cell shape (n = 40-42 cells per group from 3 independent experiments). (O) The injury response area was significantly increased between unstimulated control and 28 µM PFOS exposed larvae (*P* = 0.0306). However, there was no significant difference in response area between the 4-hour stimulated control and PFOS-exposed larvae. n = 9-15 per group. Box plot limits represent 25^th^ to 27^th^ percentile, with the midline representing the median. See Table S5 for additional statistical details.

### Assessment of global and regional neuronal network activity in PFOS-exposed larvae using CaMPARI

Given that microglia are highly responsive to their microenvironments, microglia hyperresponsiveness in PFOS-exposed larvae may be influenced by changes in neuronal communication. While evidence suggests PFOS exposure can impact developing and adult brain health, the direct effects of exposure on neuronal network activity, whole brain metabolome, and neurotransmitter release are not well understood. To understand the effect of PFOS on global brain activity *in vivo*, we assessed neuronal calcium activity by driving the fluorescent calcium sensor CaMPARI (Calcium-Modulated Photoactivatable Ratiometric Integrator) under the pan-neuronal promoter *elavl3* (*Tg(elavl3:CaMPARI(W391F+V398L))^jf9^*). CaMPARI is a permanent photoconvertible calcium sensor that undergoes allosteric chromophore modulation from green-to-red in response to ultraviolet light, but only upon simultaneous binding of free intracellular calcium^(56)^. Therefore, inactive neurons at the time of photoconversion are green, while active neurons convert to red (Figure 3A-D). In this study, free swimming larvae were subjected to 135 mW/cm^2^ of 405 nm light for 1 minute inside our Noldus behavioral unit (Figure S3A-D). We validated our CaMPARI photoconversion, imaging, and analysis pipeline by exposing larvae to pentylenetetrazol (PTZ), a GABAA receptor antagonist known to cause neuronal hyperactivity and seizures in zebrafish^(76)^ (Figure S3E-H versus 3E’-H’). Indeed, 10 mM PTZ led to a significant ratio-metric increase of neuronal intracellular calcium in the optic tectum (+22.2%) and a 16.7% increase in whole brain activity (Figure S3J). To determine whether PFOS exposure affected brain activity, we applied this validated pipeline to 3 dpf larvae exposed to 28 µM PFOS, the same concentration used for the injury model. PFOS exposure notably heightened neuronal activity in the forebrain (FB; +11.4%), optic tectum (OT; +14.1%), cerebellum (Ce; +17.9%), hindbrain (HB; +14.6%), and whole brain collectively (WB; +15.7%) (Figure 3G,I). We also examined whether lower concentrations of PFOS were able to induce differences in neuronal activity. While there were no significant differences in network activity at 3 dpf following 7 µM PFOS (Figure 3F,H), this group had notably and significantly heightened regional and global activity at 5 dpf (Figure 3K,M; FB, +7.1%; OT, +31.2%; Ce, +24.0%; HB, +15.7%; WB, +20.8%), as did the 14 μM exposed group (Figure 3L,N; FB, +14.0%; OT, +52.6%; Ce, +51.8%; HB, +34.3%; WB, +39.6%). We also assessed gross brain morphology of PFOS-exposed larvae and found that while whole brain area was slightly reduced at 3 dpf in the 7 µM (- 2.8%) and 14 µM (-2.7%) groups, brain area was unchanged by 5 dpf, suggesting that the effects of PFOS on morphology are nominal and temporary (Figure S4).

**Figure 3.**
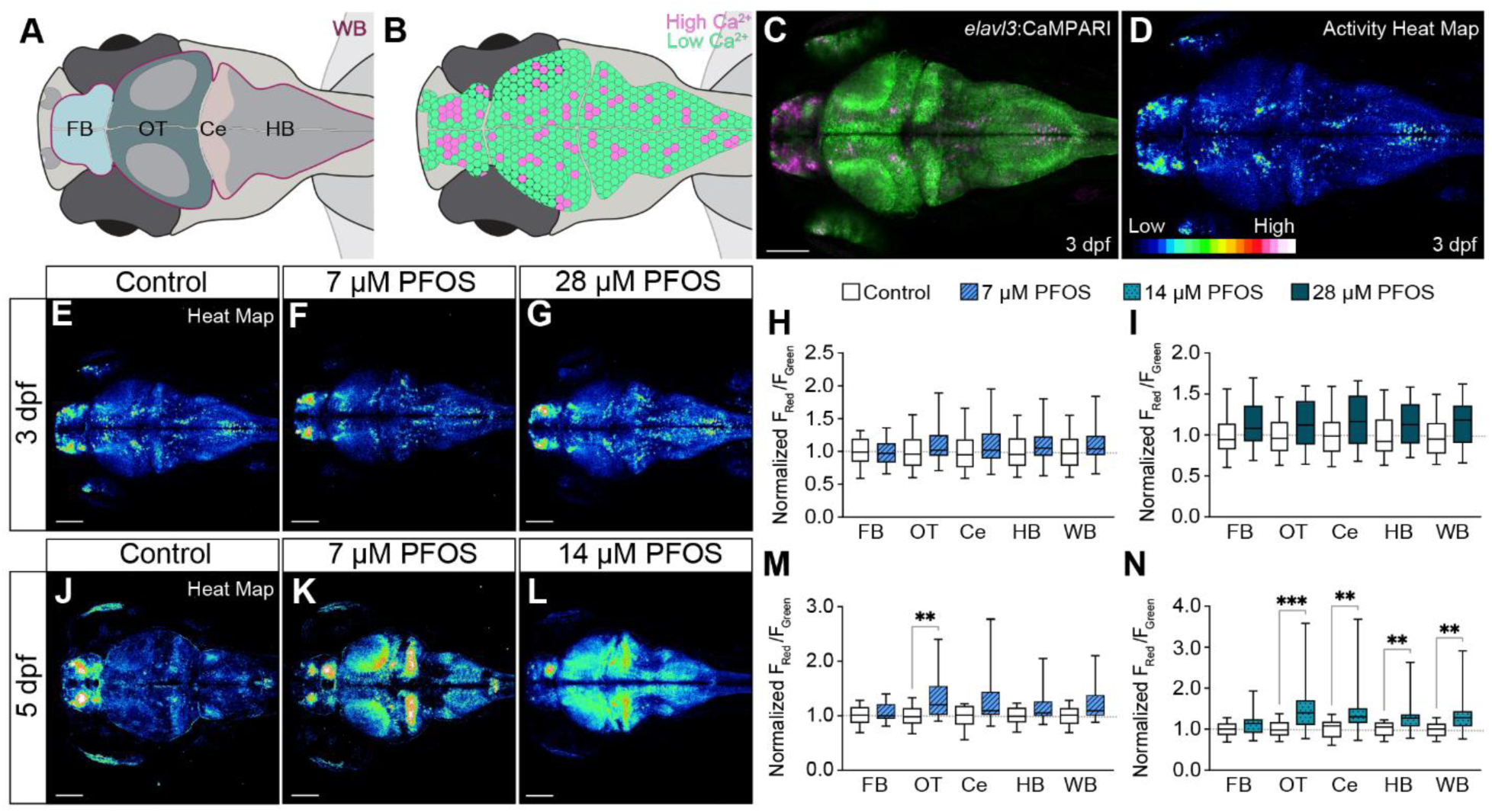
Analysis of regional and global neuronal network activity following chronic exposure to PFOS. (A) Illustrative representation of a larval zebrafish brain with anatomical regions outlined: forebrain (FB), optic tectum (OT), cerebellum (Ce), hindbrain (HB), and whole brain (WB). (B) Illustrative representation of neuron-driven CaMPARI: neurons with low intracellular calcium (low Ca^2+^) remain green following 1 minute exposure to 405 nm light, while neurons with high intracellular calcium (high Ca^2+^) are photoconverted to red. (C) Confocal micrograph of a 3 dpf larvae expressing neuron-specific CaMPARI (*Tg(elavl1:CaMPARI)*) following 1 minute photoconversion. (D) Generated high intensity LUTs heat map of the red, photoconverted channel in C depicting high activity neurons. Low to high Intracellular calcium is depicted by a blue-red-white spectrum. (E) Micrographs of active neurons at 3 dpf in control, (F) 7 µM PFOS, and (G) 28 µM PFOS larvae following 1-minute photoconversion. Neuronal activity can be quantified by determining the ratio of fluorescent intensity in the red versus green channels (F_Red_/F_Green_). (H) 7 µM PFOS does not result in a regional or global (WB) change in neuron activity in 3 dpf larvae. (J) Brain activity was also determined in 5 dpf larvae exposed to control, (K) 7 µM PFOS, or (L) 14 µM PFOS. At 5 dpf (M) 7 µM PFOS-exposed larvae have significant increases in brain activity in the OT (*P* = 0.0058) and near significance globally (*P* = 0.0788). (N) Larvae exposed to 14 µM PFOS also had increases in the OT (*P* = 0.0005), Ce (*P* = 0.0010), HB (*P* = 0.0074), and globally (*P* = 0.0025). Confocal micrographs at 10x magnification. n = 21-23 fish per group. Box plot limits represent 25^th^ to 27^th^ percentile, with the midline representing the median. See Table S5 for additional statistical details.

To better understand the neurochemical changes in the PFOS exposed brain, we performed an untargeted metabolome wide association study (MWAS) on isolated heads of 3 dpf control and 28 µM PFOS exposed larvae. Several features were significantly upregulated or downregulated between the control and exposed groups (Figure S5). Significantly different metabolites from the MWAS were further analyzed for pathway enrichment using MetaboAnalystR^(68)^. Several of the significantly enriched pathways, including glutamate metabolism, aspartate and asparagine metabolism, tyrosine metabolism, and phosphoinositide metabolism, are involved in neuronal excitation, catecholamine synthesis, and neurotransmitter release, uptake, and recycling (Figure S5C; Table S4).

We next sought to determine whether the neurochemical imbalances were predictors of abnormal behavioral activity. To first determine whether our exposure paradigm replicates previously reported behavioral hyperactivity, we exposed embryos to either 7 μM, 14 μM, or 28 μM PFOS at 4 hpf, and performed light/dark behavioral assays at 3 dpf, 4 dpf, and 5 dpf (Figure S6A,B). Consistent with previous locomotor assays at 6 dpf^(48)^, PFOS-exposed larvae were significantly more responsive to light changes, designated by elevated distance traveled within the well at 3 dpf following 28 μM exposure (Figure S6E-H) and 5 dpf following 7 μM and 14 μM exposure (Figure S6N-R). In addition to distance traveled, behavior videos were analyzed to assess the frequency of center avoidance within the wells of a 24-well plate. Similar to the mammalian open-field test, center avoidance in a well is an indication of zebrafish anxiety-like behavior, while willingness to cross the center suggests a reduced anxiety-like behavior^(77)^. As such, we examined the time each larva spent along the well’s edge (anxiety-like) versus center (exploratory) throughout each light/dark behavioral assay (Figure 4A,B). PFOS exposure resulted in notable and significant reductions in time spent in the center in all dosed groups at 3 and 5 dpf (Figure 4C-E). Interestingly, while the 7 µM PFOS group were not significantly more active during the light on cycle at 5 dpf (Figure S6P), they only exhibited center-avoidance during the light on cycle (Figure S7G,H). Likewise, while 14 μM PFOS-exposed larvae exhibited significant center-avoidance behavior at 3 dpf, they did not have increased swim activity at 3 dpf (Figure S6H). Further, the 14 µM PFOS group exhibited center-avoidance behavior in both the light and dark cycles at 5 dpf (Figure S7I,J), though were only significantly more active when the light was on, and not when the light was off (Figure S6P,Q). This suggests that anxiety-like behaviors are separable from swim hyperactivity and supports the need to more thoroughly understand the independent behavioral changes associated with PFOS exposure. Since anxiety-, fear-, and aversive-like behaviors are regulated by the medial habenula in zebrafish^(78, 79)^, we assessed neuron activity at the habenula using CaMPARI. PFOS exposure did not result in a difference in habenular activity at 3 dpf, and only 14 µM PFOS-exposed larvae showed a significant difference in habenular activity at 5 dpf (Figure S8).

**Figure 4.**
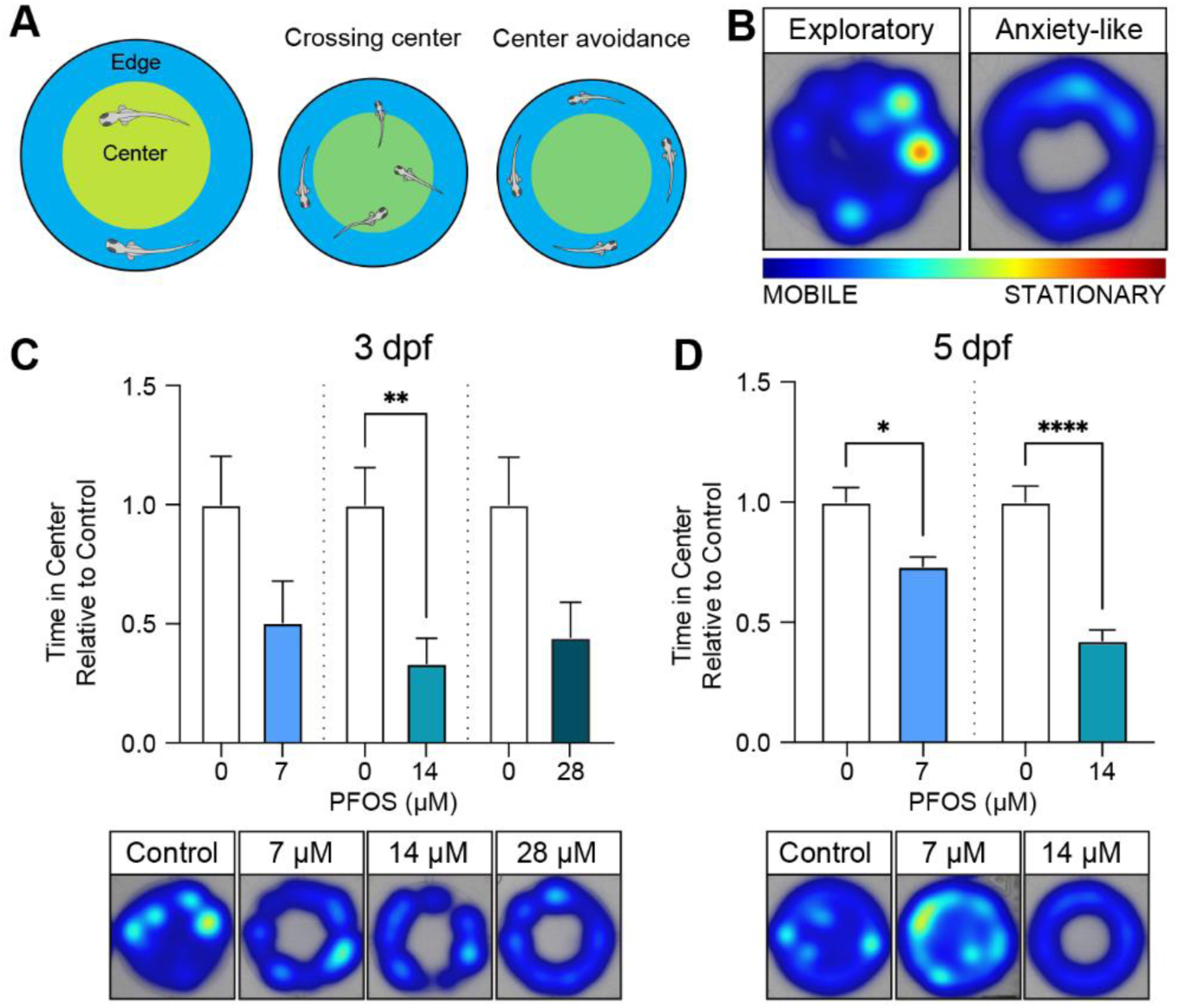
Assessing anxiety-like swim behavior in PFOS-exposed larvae. (A) Larval swim behavior during a 30-minute light/dark behavioral assay was used to determine time spent in the well’s center (crossing center; less anxious) versus time spent along the well’s edge (center avoidance; more anxious). (B) Heat maps were generated to indicate mobile (blue) versus stationary (red) swim activity within the well, as well as the zones traversed within the well. (C) At 3 dpf, larvae exposed to PFOS spent less time in the center relative to the corresponding control group (7 µM, -49.5%, *P =* 0.2994; 14 µM, -66.5%, *P* = 0.0021;28 µM, -55.6%, *P* = 0.6682). (D) At 5 dpf, both 7 µM PFOS (-26.7%; *P* = 0.0288) and 14 µM PFOS (-57.5%; *P* < 0.0001) spent significantly less time in the center of the well. n = 76-101 fish per group. Error bars represent SEM. See Table S5 for additional statistical details.

### Evaluation of light/dark response in microglia mutants exposed to PFOS

Neuron-microglia bi-directional signaling enables the reciprocal regulation of microglia and neuronal functions, including microglial modulation of neuronal activity and circuit refinement^(35, 80)^. Considering the observed shift in microglial state following PFOS exposure, we sought to determine whether or not PFOS-induced microglial dysfunction during early larval development influenced neuronal excitation and behavior. To do this, we exposed PFOS to zebrafish with a null mutation for *irf8* (*irf8^st96/st96^*), a gene required for macrophage formation during primitive and transient definitive hematopoiesis^(60)^. *Irf8* mutant larvae lack the earliest embryonic-derived macrophage populations, including microglia^(60)^. Control-treated *irf8* mutants had a slight reduction in swim behavior throughout the recording (Figure 5A; gray versus yellow traces), though were significantly less reactive to the light-to-dark transitions than control-treated wildtype larvae (Figure 5B; 96.8 ± 8.3 mm moved versus 71.5 ± 6.8 mm moved). At 5 dpf, both wildtype and *irf8* mutant larvae exposed to 8 µM PFOS were significantly more active than control-treated larvae (Figure 5A; Figure S9E-G). However, unlike the control-treated *irf8* mutants, PFOS-exposed mutants did not display a trend toward reduced swim activity during the light/dark behavioral assay (Figure 5A; blue versus red traces). PFOS-exposed mutants also exhibited the most dramatic behavioral response to the light-to-dark transitions, especially during the second dark cycle (Figure 5B; “Minute 15”). In addition, while wildtype control larvae and wildtype PFOS-exposed larvae had a 14.9% and 23.8% reduction in swim activity by the end of the assay (“Minute 30”), respectively, PFOS-exposed *irf8* mutants had a 47.6% reduction in activity (Figure 5B). These data suggest that microglia may attempt to modulate the PFOS-exposed brain, such that the absence of microglia may further promote PFOS-induced behavioral reactivity and altered patterns of activity post-dark transitions. Heightened reactivity may also result in behaviors that impair swim ability, such as seizing or convulsions. PFOS exposure also results in increases in neuronal-driven CaMPARI in microglia-deficient larvae, but this was not significant between genotypes (Figure 5C).

**Figure 5.**
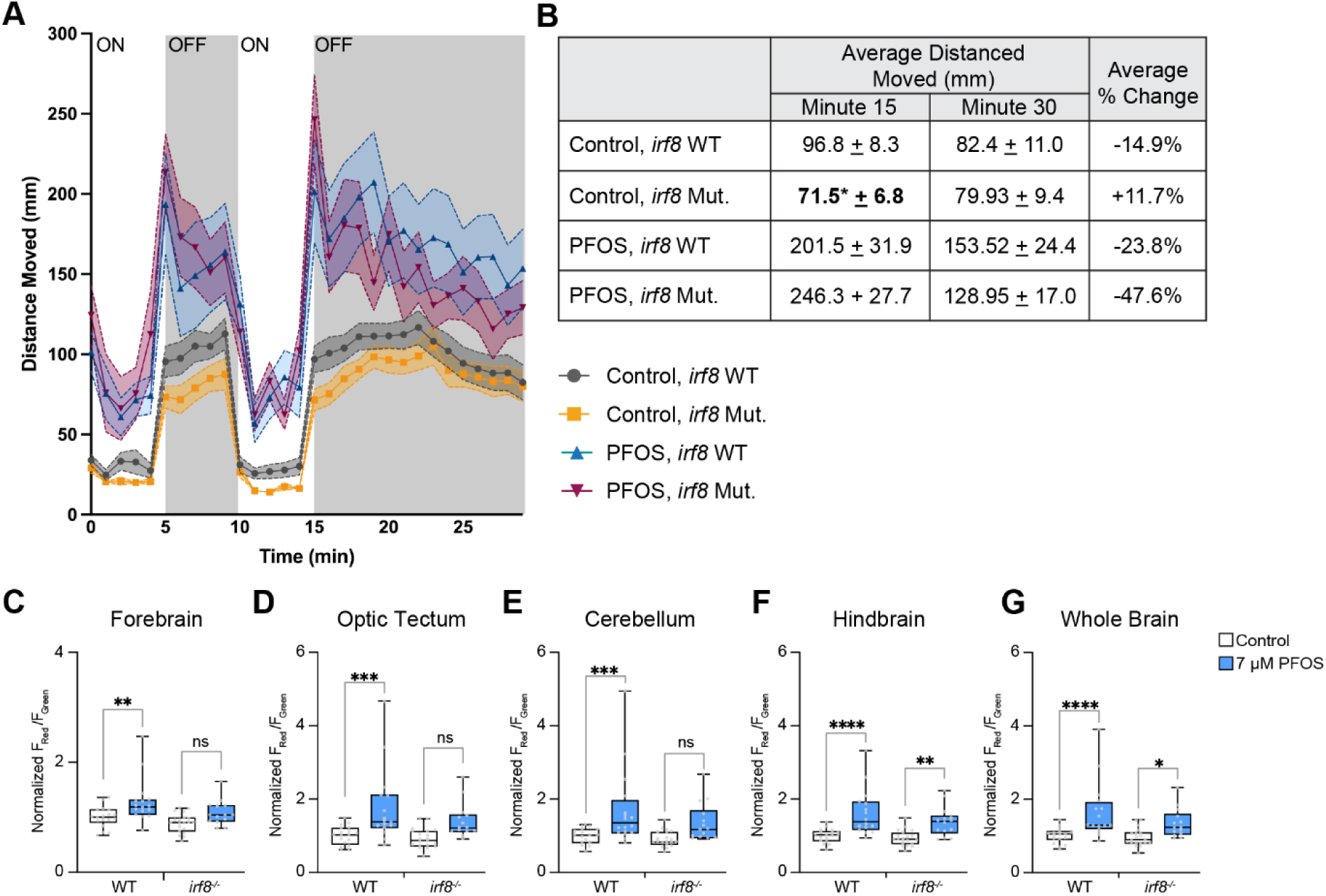
Light/dark behavioral assay and neuronal activity in microglia mutants exposed to PFOS. Wildtype and microglia-deficient *irf8^st96/st96^* larvae dosed with either control or 7 µM PFOS were assessed for potential changes in swim behavior and neuronal activity. (A) Light/dark behavioral assay of 5 dpf wildtype or *irf8* mutant control or PFOS-exposed larvae. (B) Assessing the distance moved following the first minute of the second light “off” cycle (i.e., minute 15 of the assay), control-treated *irf8* mutant larvae responded significantly less than wildtype controls (*P* = 0.0237). While not significant, PFOS-exposed *irf8* mutants had a heightened response to the first minute of the dark cycle compared to PFOS-exposed wildtype larvae. PFOS mutants also had twice the percent recovery in swim activity by the last minute of the assay (i.e., minute 15 versus minute 30). (C-G) PFOS-exposed macrophage mutants did not show a significant change in regional or global neuronal activity compared to PFOS-exposed wildtype larvae. (C) Forebrain, control-treated wildtype versus PFOS-treated wildtype (*P* = 0.0060); control-treated mutant versus PFOS-treated mutant (*P* = 0.1176). (D) Optic tectum, control-treated wildtype versus PFOS-treated wildtype (*P* = 0.0008); control-treated mutant versus PFOS-treated mutant (*P* = 0.0867). (E) Cerebellum, control-treated wildtype versus PFOS-treated wildtype (*P* = 0.0020); control-treated mutant versus PFOS-treated mutant (*P* = 0.1333). (F) Hindbrain, control-treated wildtype versus PFOS-treated wildtype (*P* < 0.0001); control-treated mutant versus PFOS-treated mutant (*P* = 0.0100). (G) Whole brain, control-treated wildtype versus PFOS-treated wildtype (*P* = 0.0001); control-treated mutant versus PFOS-treated mutant (*P* = 0.0349). There were no significant differences in regional or global PFOS-induced neuronal hyperactivity between the wildtype or mutant larvae expressing *Tg*(*elavl3:CaMPARI*). n = 14-22 per group for behavior; n = 13-16 per group for neuroimaging. Box plot limits represent 25^th^ to 27^th^ percentile, with the midline representing the median. See Table S5 for additional statistical details.

### Microglia response to brain injury following optogenetic modulation of neuronal activity

Given the bi-directional communication between microglia and neurons and that microglial state can be modulated by neural activity, we asked whether elevated neural activity alone was sufficient to modify microglial responses to injury. To test this hypothesis, 3 dpf larvae were treated with 5 mM PTZ to increase neuronal activity, then were injured in the right telencephalon as described. PTZ-induced neuronal hyperactivity significantly increased the microglia response at 4 hpi, similar to that of PFOS-exposed larvae (Figure S10). We next asked whether inhibiting neuronal activity in PFOS-exposed larvae using optogenetics could normalize the observed hyperresponsive microglial phenotype. Larvae expressing *Tg(elavl3:Gal4;cryaa:RFP;UAS:eNpHR3.0;mpeg1:EGFP),* which have pan-neuronal expression of halorhodopsin as well as GFP-labeled macrophages, were exposed to either a control solution or 28 μM PFOS at 4 hpf. After brain injury at 3 dpf, larvae were either left unstimulated or stimulated for 4 hours with 570 nm light to silence neuronal activity. Microglia of unstimulated PFOS-exposed larvae were 34.5% more responsive to injury (Figure 6E,F,I). Conversely, neuronal hyperpolarization rescued microglia hyperresponsiveness, resulting in a 6.6% decrease in microglia response in stimulated PFOS-exposed larvae compared to stimulated controls (Figure 6G,H,I), further supporting that pollutant-induced changes in the neural signaling environment significantly influences microglia behavior, independent of cell death or inflammation.

**Figure 6.**
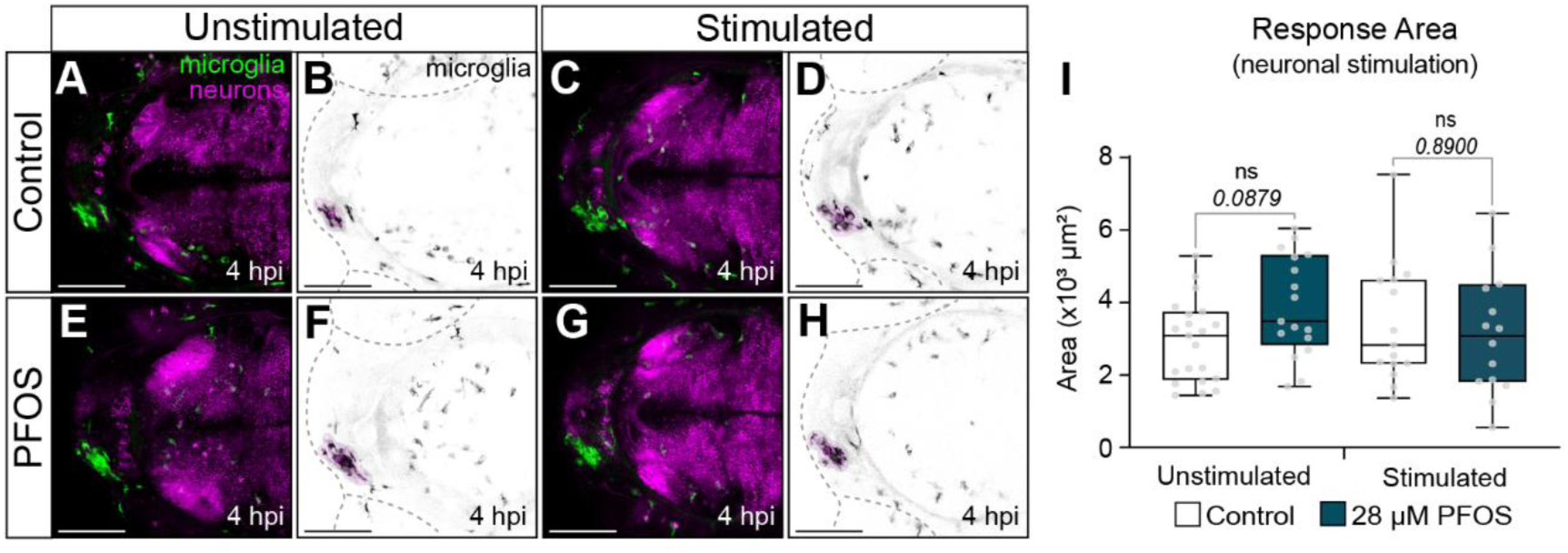
Microglia response to brain injury following optogenetic silencing of neurons in PFOS-exposed larvae. Larvae expressing *Tg(elavl3:Gal4;cryaa:RFP;UAS:eNpHR3.0;mpeg1:EGFP)* were dosed with (A-D) control or (E-H) 28 µM PFOS and injured at the right telencephalon at 3 dpf and imaged 4 hpi. (E&F) PFOS-exposed unstimulated larvae had a 33.5% increase in microglia response compared to controls (*P* = 0.0879). (C&D) Optogenetic stimulation of halorhodopsin with 570 nm light for 4 hpi did not affect microglia response to injury in control larvae (6.6% decrease). (G&H) However, neuronal silencing in PFOS-exposed larvae normalized the microglia response, such that the response was equivalent to the stimulated controls. (I) Quantification of microglia response area. n = 14-20 per group. Box plot limits represent 25^th^ to 27^th^ percentile, with the midline representing the median. See Table S5 for additional statistical details.

### Microglia response to brain injury and neuronal network activity following chronic exposure to PFOA, a known immunotoxic congener

To further assess if microglia hyperresponsiveness following PFOS exposure is a result of neuronal hyperactivity, we exposed larval zebrafish to a structurally similar PFAS congener, perfluorooctanoic acid (PFOA). Whereas PFOS has a sulfonic acid head group, PFOA is an 8-carbon PFAS with a carboxylic acid head group. Following the same exposure paradigm used for PFOS exposure (Figure 1A), zebrafish embryos were dosed with either a control solution (0.1% DMSO) or 64 µM PFOA at 4 hpf. Exposure to 64 µM PFOA did not result in differing swim behavior during light/dark behavioral assays at 5 dpf (Figure S11A-D). While PFOA-exposed larvae had shorter body length, there were no observable gross morphological effects on the spine that would affect swimming ability (Figure S12). Using neuronally-driven CaMPARI, PFOA exposure did not result in any regional or global differences in neuronal network activity in 5 dpf larvae (Figure 7A-C), unlike the PFOS-exposed groups at this timepoint (Figure 3). Given that PFOA did not result in neuronal hyperactivity, we asked whether PFOA exposure affected microglial responses to brain injury. Indeed, PFOA exposure did not result in differing microglial responses to brain injury (Figure 7D-F). Together, these data further support that neuronal hyperactivity is a key driver of microglia response, and that structurally similar PFAS congeners can have distinct impacts on the CNS.

**Figure 7.**
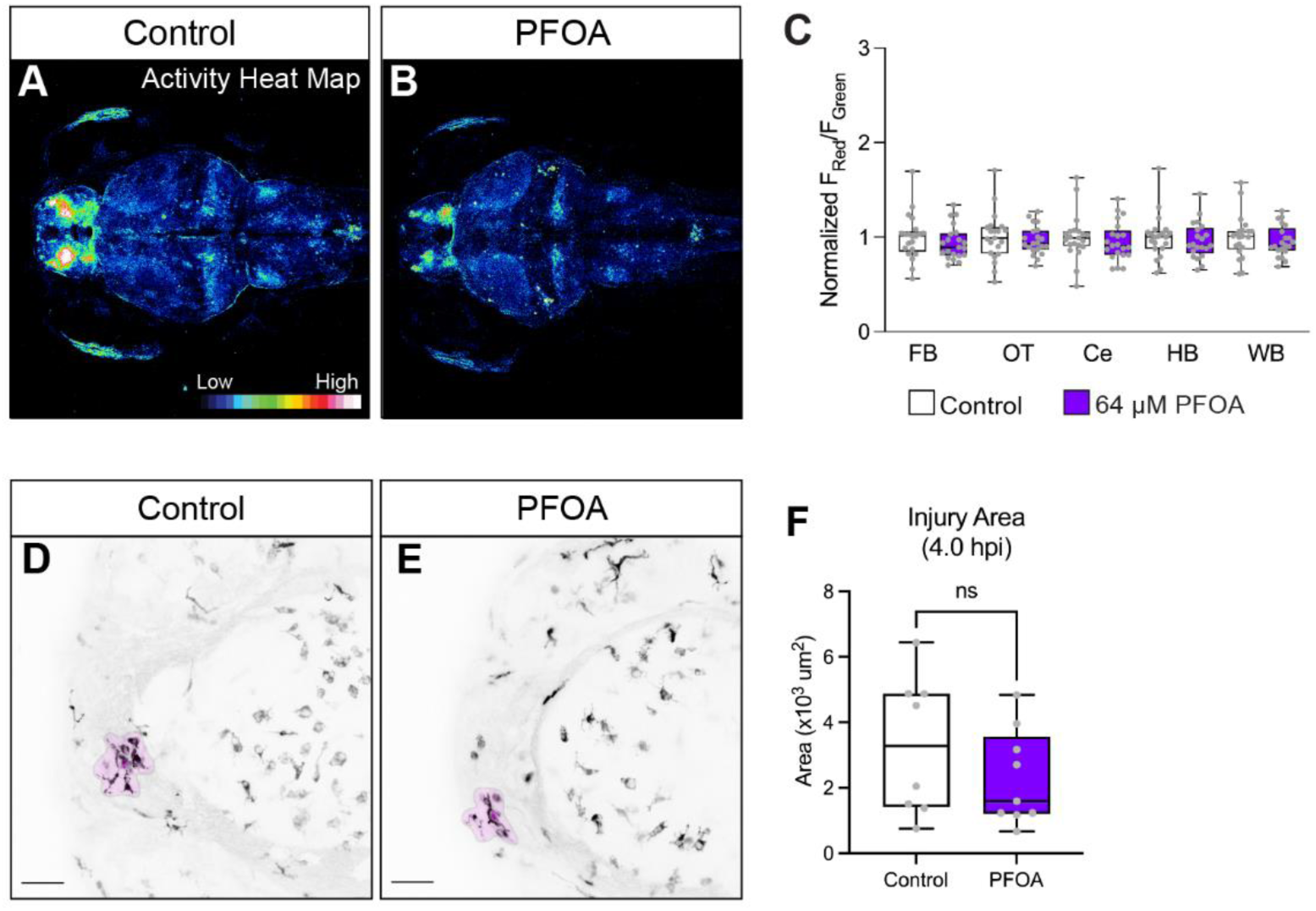
Neuronal activity and microglia response to brain injury following exposure to PFOA, a known immunotoxic congener. PFOA is an 8-carbon PFAS compound with a carboxylate head group. To observe the effects of PFOA on neuronal activity, we again used larvae of the Tg(*elavl3:CaMPARI*) background. (A) At 5 dpf, control and (B) 64 µM PFOA-exposed larvae show no significant changes in regional or global neuronal activity, quantified in (C). (D) Larvae expressing Tg*(mpeg1:EGFP)* were also dosed with control solution or (E) 64 µM PFOA to determine if this non-excitatory PFAS compound resulted in microglia response differences at 3dpf. (F) At 4 hpi, 64 µM PFOA-exposed larvae did not have a significant change in microglia response to injury compared to sibling controls. n = 8-9 per group. Box plot limits represent 25^th^ to 27^th^ percentile, with the midline representing the median. See Table S5 for additional statistical details.

## DISCUSSION

For more than two decades, researchers have revealed the many biologic, ecologic, and environmental ramifications of PFOS toxicity, as well as the toxicity profiles for a subset of other ‘forever chemical’ congeners. Here, we used a combination of *in vivo* imaging of cellular behavior, functional neuroimaging, optogenetic modulation, and behavioral assays to address the impact of PFOS exposure on microglia-neuron interactions and activity in larval zebrafish. Our data demonstrated that developmental PFOS exposure modulated microglia state and induced hyperactivity in response to injury independent of cell death or inflammation. Since the homeostatic microglia membrane potential is largely maintained by chloride channel currents^(75)^, we asked whether activating the optogenetic chloride channel halorhodopsin (eNpHR3.0) could rescue microglia hyperresponsiveness. Indeed, electrical modulation of microglia was sufficient to revert microglia from an amoeboid to a homeostatic morphology and normalize their responses to injury, suggesting electrical dysfunction as a previously underappreciated pathway worth interrogating when studying microglial state and immunotoxicity. PFOS-exposed larvae also exhibited global and regional increases in neuronal activity and anxiety-like behaviors, a previously unidentified neurodevelopmental phenotype of PFOS exposure in zebrafish, with only nominal and temporary impacts on regional brain morphology. We demonstrated that PFOS-induced neuronal hyperactivity was a key mediator of microglia reactivity and that optogenetic silencing of neurons was sufficient to normalize microglia responses to injury. Further, exposure to PFOA did not result in neuronal hyperactivity nor microglia hyperresponsiveness. Together, this study provides the first detailed account of the effects of PFOS exposure on the developing brain *in vivo* and adds neuronal hyperactivity as an important endpoint to assess when studying the impact of toxicant exposures on microglia function.

Zebrafish have long been considered excellent models to study innate immune development and function^(81, 82)^, including to understand neuro-immune interactions^(83)^. While the immunotoxic impact of PFOS on adaptive immunity has been well documented^(84)^, the effects on the innate immune arm are less understood and, at times, contradictory. Studies showing either innate immune activation or suppression following PFOS exposure are likely attributed to varying exposure paradigms, PFOS concentration, animal model, age, or cell line used^(24, 26, 28, 85, 86)^. The innate immune system is highly sensitive to both endogenous and xenobiotic stimuli and reacts rapidly, creating a signaling cascade that informs all downstream immune functions^(87)^. Macrophages in particular have the ability to dynamically polarize from a homeostatic state to be pro- or anti-inflammatory depending on the environmental needs. As antigen-presenting cells, macrophages also have the vital role of instructing adaptive immune cells on their responses^(87)^. The dependency of the adaptive immune system on the innate emphasizes the need to clarify the immunotoxic mechanisms of pollutants like PFOS.

This work constitutes the first *in vivo* analysis of microglia, the resident immune population of the CNS, following PFAS exposure. Previous *in vitro* studies using immortalized microglia cell lines suggested PFOS exposure decreased microglial viability, mitochondrial stability, and increased ROS production *in vitro* in a concentration-dependent manner^(88, 89)^. We found that PFOS exposure altered the microglia state, demonstrated by their morphological transition from ramified to amoeboid shaped, as well as by the upregulation of the microglia activation gene *p2ry12,* a purinergic receptor that responds to ATP. It was previously demonstrated that microglia response to injury was mediated by glutamate-evoked calcium waves and ATP release from the injury site^(73)^; therefore, ATP released by the injury site and/or through high neuronal activity could be contributing to the microglial hyperresponsiveness in PFOS-exposed larvae. It is still unclear to what extent PFOS is acting directly on the microglia or indirectly through neuronal activation. Microglia activation and injury responses were attenuated by both microglia and neuronal optogenetic silencing, suggesting that the neuronal environment has a significant influence on the microglia. Of note, while larvae with neuron-driven halorhodopsin had a 34.5% higher microglial response to injury following PFOS exposure, the response was not as significant as it was in other genetic backgrounds (*P* = 0.0879). This is likely due to the fact that optogenetic channels and pumps are highly sensitive, and while we reduced potential exposure to ambient light, leakiness of these opsins may contribute to low level ion exchange. More work on the baseline permeability of optogenetic channels in a controlled and cell-specific context is needed.

While we are providing the first demonstration of microglial dysfunction in response to toxicant exposure, a shift away from a homeostatic microglial state has been well documented in disease models of various neuropathological states. In addition, pharmacological manipulation of microglia state or prevention of microglia depletion was shown to lead to cognitive and functional improvements in some neurological disease models^(37, 38, 90)^. However, it is worth noting that modulation of microglia state is not inherently pathological. For example, microglia depletion following stroke significantly increased infarct size, neuronal cell death, and caused calcium overload^(91)^. Microglia depletion during acute seizures also exacerbated excitotoxicity and seizure sensitivity^(92)^. Additionally, the regional specificity of microglia state in a mouse model of chronic stress was considered a protective and/or adaptive response^(93)^. Lastly, it is important to note that the dosing paradigm used in this study has been shown to impact other organ systems, such as the pancreas^(61)^. Therefore, one cannot dismiss the potential for systemic effects contributing to the neurochemical imbalances or altered microglial state. While our timelapse imaging videos, cell death and inflammation data, and microglia quantifications do not suggest macrophages are recruited from the periphery to the brain injury site, more extensive analysis using cell lineage tracing from the periphery would need to be performed to confirm this. However, the potential for the periphery to impact brain health further emphasizes the benefit of understanding systems-level interactions when studying PFAS exposure.

Due to the developmental and homeostatic roles microglia have on maintaining the excitatory/inhibitory balance^(94)^, as well as the situation-specific adverse effects that changes in microglia state may have on the CNS, we asked whether modulation of microglial state following PFOS exposure was contributing to the behavioral or neuronal hyperactivity. Using microglia-deficient zebrafish with a mutation in the gene *irf8*, we found that control-treated *irf8* mutants had a slight but consistent reduction in swim behavior that was not seen in the PFOS-treated mutants. Additionally, PFOS-exposed mutants had the greatest response to the light-dark transition, suggesting that microglia may actually attempt to repress PFOS-induced behavioral hyperactivity. While microglia loss did not affect baseline neuronal calcium signaling, it is worth noting that microglia loss is distinct from microglial dysfunction, and thus does not rule out the contribution of microglia to neuronal hyperactivity in this context. For example, signals derived from dysfunctional microglial could impact neuronal firing and perhaps swim behavioral responses, which would not be recapitulated in a model of microglial loss. In addition, microglia increasingly accumulate in synaptic regions between 7 and 28 dpf in zebrafish^(95)^, suggesting that any significant death-dependent or activity-dependent neuronal pruning may occur developmentally later than the timepoints investigated in this study. Lastly, the concentrations of PFOS used in this study might increase neuronal activity too substantially or even irreversibly, such that electrical modulation or loss of microglia is not sufficient to influence the activity in a measurable way.

To our knowledge, this is the first report of *in vivo* neuronal hyperactivity caused by embryonic PFOS exposure. We show that embryonic PFOS exposure resulted in higher intracellular calcium concentrations across multiple brain regions at various concentrations (7 µM, 14 µM, and 28 µM) and time points (3 dpf & 5 dpf). Both gestational and adult PFOS exposures have been shown to impact calcium dependent signaling molecules important for memory, including Ca^2+^/calmodulin dependent kinase II (CaMKII) and cAMP response element-binding protein (CREB) in rats^(96)^, suggesting that disrupted calcium signaling is an important mediator of PFOS-induced neurotoxicity. *In vitro* studies also linked PFOS toxicity to disrupted calcium homeostasis^(51, 97)^. While the underlying mechanisms remain unknown, possible explanations include increased influx through activation of L-type Ca^2+^ channels^(97)^ and release of intracellular calcium stores through interaction with ryanodine and inositol 1,4,5-trisphosphate receptors^(51)^. Intracellular calcium excess in neurons promotes excitotoxicity and can cause brain damage leading to various neurological and neurodegenerative disorders^(98, 99)^, emphasizing the need to further understand the influence of PFAS on neuronal function. PFAS compounds have also been shown to activate peroxisome proliferator-activated receptors (PPARs) *in vitro* as well as in zebrafish^(100, 101)^. PPAR-γ in particular is expressed in neurons, microglia, and astrocytes and mediates inflammatory responses in the CNS^(102)^. Interrogating PPAR activation in the CNS following PFOS exposure could also provide important information about the specific pathways impacted by exposure.

The susceptibility of the developing human brain to PFOS remains a contentious point due to conflicting data. While some studies have demonstrated significant correlations between developmental PFOS exposure and ADHD incidence^(40, 41)^, others found no such relationship^(42–45, 103)^. In the mouse brain, PFOS concentrations have been shown to increase over time and lead to tonic convulsions, despite a lack of morphological phenotypes^(104)^. The heightened regional and global network activity demonstrated in this study warrants an assessment of convulsive phenotypes and seizure activity. While outside the scope of this study, determining the threshold and timescale at which neuronal hyperactivity increases incidence of convulsion would be interesting to pursue.

Similar to previous reports^(48–50)^, we observed hyperactive swim behavior in PFOS-exposed larvae during light/dark behavioral assays. However, we are the first to report PFOS-induced hyperactivity during the light and dark phases prior to 6 dpf. Hyperactive behavioral changes corresponded well with increased intracellular neuronal calcium concentrations. Of note, increased anxiety-like behaviors were separable from increased swim activity at 3 dpf and 5 dpf and was also specific to light or dark cycles depending on PFOS concentration. This suggests that there may be further disruptions in neuronal communication beyond just increased calcium concentrations influencing swim activity. In-depth regional brain activity analyses, possibly through the integration of developed analytical pipelines^(105)^, could provide further insight into how PFOS induced neural activation is linked to behavioral abnormalities in the larval zebrafish.

We identify increased anxiety-like behaviors as a novel phenotype of PFOS neurotoxicity in the larval zebrafish using an adapted open-field test model^(106–108)^. Anxiety is an associated symptom of several neurobehavioral disorders linked to PFAS exposures, including autism spectrum disorder and ADHD^(45, 109–111)^. PFOS-induced center avoidance has been observed in mice exposed during adulthood^(112)^, but developmental anxiety-like behavior has not been previously reported. Of note, previous published research reporting PFOS-induced larval hyperactivity utilized 96-well plates for higher throughput^(48, 50)^; however, smaller well sizes may not be conducive to conducting anxiety-like behavioral analyses. We therefore conducted behavioral experiments using 24-well plates, which allowed us to define regions within the well to quantify where fish spent their time swimming. Because dysregulation of habenular activity has been associated with increased anxiety, depression, and fear across species, including the zebrafish^(78, 79, 113)^, we assessed whether neuronal calcium was affected in the habenula following PFOS exposure. While only the 14 µM PFOS group showed neuronal dysregulation at the habenula, all groups displayed anxiety-like behaviors. This suggests that the habenula may be but one region dictating anxiety-like responses in larval zebrafish or that temporal differences in habenular activity, not captured using CaMPARI, contribute to anxiety-like responses.

Not only does this study provide the first *in vivo* assessment of neuronal calcium activity following PFOS exposure, but it is also the first to evaluate PFOA exposure in this context. The chemical structure of PFOA is similar to PFOS, differing only by the presence of a carboxylic acid head group rather than sulfonic acid, respectively. PFOA has been shown to cause significant health effects, including immunotoxicity, elevated cholesterol, dysregulated liver metabolism, kidney dysfunction, thyroid disease, among many others^(114)^. Previous reports using larval zebrafish have found that PFOA concentrations ranging from 4.4 µM to 80 µM did not result in a significant difference in 6 dpf swim behavior during the light/dark behavioral assays^(48)^. At the concentrations and timepoints tested here, PFOA exposure did not result in heightened larval swim behavior or neuronal network activity. However, others have reported that 5 dpf larvae exposed to 1 µM PFOA^(115)^ as well as 14 dpf larvae exposed to 2 µM PFOA^(50)^ did result in swim hyperactivity. In addition, while neonatal exposure to PFOA in mice has been shown to result in abnormal expression of proteins important for brain growth and neuron function^(46)^, PFOA exposure of rat primary cortical neurons did not affect spontaneous neuronal activity or burst duration^(116)^. Further analysis is needed to clarify the extent to which PFOA, and other shorter chain length or carboxylic group PFAS, could be neurotoxic or contribute to neural dysregulation. Nevertheless, that PFOA-exposed larvae had normal microglia responses to injury, compared to those exposed to PFOS, further highlights the role of neuronal activity on microglia function.

This study provides the first *in vivo* analysis of how developmental PFOS exposure disrupts larval brain health and function. It also highlights the relevance of understanding pollution-induced effects on innate immune cells, in both a non-canonical developmental and homeostatic context, as well as considering the long-term consequences of potential antigen-presentation dysfunction. In summary, the complexity of neural cell communication, especially during the sensitive period of brain development, emphasizes the importance of studying pollutant exposure in non-isolated systems.

## Supporting information

Supplemental Video 1_Control

Supplemental Video 2_PFOS

## ACKNOWLEDGMENTS

We thank Dr. William Talbot for sharing their *irf8^st96/st96^* mutant line with us. We also thank Rachel Cyr for excellent care and oversight of our zebrafish facility. Thank you to other members of the Plavicki lab for their discussions in input during various stages of this study. Lastly, thank you to the undergraduate trainees that were involved in the early coordination of this work, Rekha Dhillon-Richardson and Ratna Patel. This work was supported by a NIEHS Outstanding New Environmental Scientist (ONES) award, and cardiopulmonary vascular COBRE Phase II (2PG20GM103652) awarded to J.S.P. In addition, this work was supported by the Ruth L. Kirschstein Predoctoral Individual National Research Service Award (NRSA; F31HL156460) by the NHLBI awarded to S.E.P. S.E.P and N.R.M. were previously supported by the Brown University Environmental Pathology Training Grant (T32ES007272-26) from NIEHS. The Thermo LC-Orbitrap MS was partially funded by NSF Major Research Instrumentation (MRI) award CBET-1919870 to K.P. (PI) and J.S.P (Co-I).

## Supplemental Material

### Supplemental Tables

**Table S1.** Survival rate of zebrafish larvae exposed to PFOS over time.

| Age | Control | 14 μM PFOS | 28 μM PFOS | 56 μM PFOS |
| --- | --- | --- | --- | --- |
| 24 hpf | 97.93 ± 2.94 | 97.67 ± 1.75 | 98.15 ± 1.73 | 97.22 ± 2.84 |
| 48 hpf | 97.74 ± 2.76 | 97.14 ± 1.21 | 97.09 ± 2.10 | 2.49 ± 2.84 |
| 72 hpf | 97.43 ± 3.25 | 96.18 ± 1.81 | 95.62 ± 2.43 | 81.54 ± 13.23 |
| 96 hpf | 96.41 ± 5.82 | 95.72 ± 1.20 | 50.65 ± 10.06* | 25.00 ± 9.89** |
| 120 hpf | 95.93 ± 5.85 | 89.14 ± 3.65 | 7.73 ± 3.03**** | 5.33 ± 2.67**** |

Percent survival of larvae aged 24 hpf to 120 hpf exposed to control solution, 14 μM PFOS, 28 μM PFOS, or 56 μM PFOS. Statistics compared each timepoint to the 24 hpf timepoint of the same treatment group using unpaired t-tests. ± denotes standard deviation. \* $P$ <0.05; \*\*\* $P$ <0.001; \*\*\*\* $P$ <0.0001.

**Table S2.**
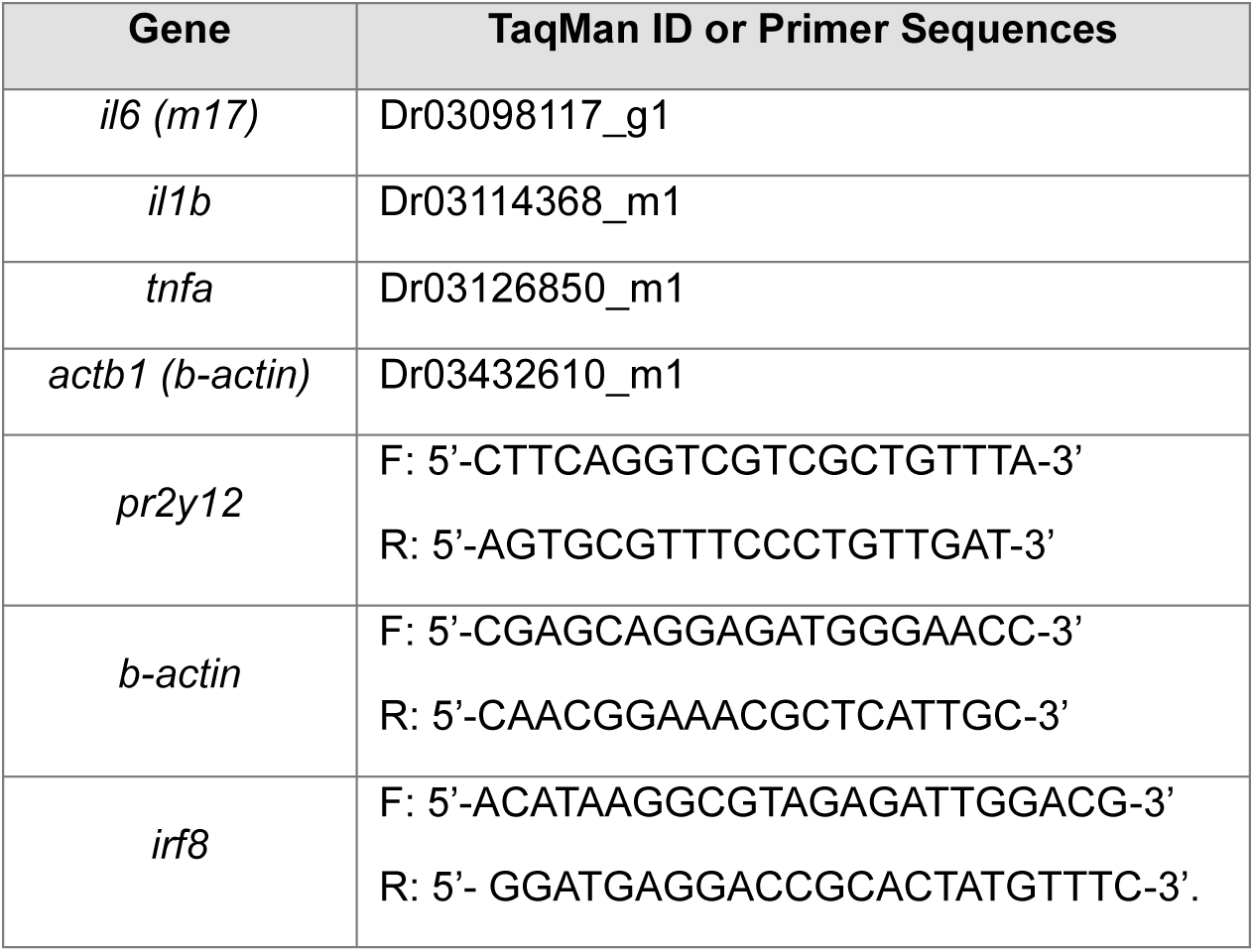
Primers used for qRT-PCR and genotyping.

**Table S3.** PFOS body burden in zebrafish larvae.

| Chemical | Formula | MW<br>(g/mol) | Control | | 28 μM PFOS | | 56 μM PFOS | |
| --- | --- | --- | --- | --- | --- | --- | --- | --- |
|  |  |  | 48 hpf | 72 hpf | 48 hpf | 72 hpf | 48 hpf | 72 hpf |
| PFOS | C<sub>8</sub>HF<sub>17</sub>O<sub>3</sub>S | 500.13 | 0.12 ± 0.17 | 0.12 ± 0.16 | 32.22 ± 2.57 | **70.46 ± 2.72\*\*\*\*** | 46.04 ± 3.75 | **109.15 ± 5.99\*\*\*\*** |
| PFNS | C<sub>9</sub>HF<sub>19</sub>O<sub>3</sub>S | 550.14 | 0.01 ± 0.02 | 0.00 ± 0.00 | 0.19 ± 0.05 | **0.39 ± 0.07\*\*\*\*** | 0.23 ± 0.04 | **0.69 ± 0.20\*\*\*** |
| PFHpS | C<sub>7</sub>HF<sub>15</sub>O<sub>3</sub>S | 450.12 | 0.00 ± 0.00 | 0.00 ± 0.00 | 0.35 ± 0.17 | 0.45 ± 0.13 | 0.40 ± 0.17 | **0.55 ± 0.11\*** |
| PFHxS | C<sub>6</sub>HF<sub>13</sub>O<sub>3</sub>S | 400.12 | 0.006 ± 0.01 | 0.008 ± 0.01 | 0.008 ± 0.01 | 0.007 ± 0.01 | 0.008 ± 0.01 | 0.008 ± 0.01 |

PFOS body burden (ng/embryo) in 48 hpf and 72 hpf larvae dosed with control solution, 28 μM PFOS, or 56 μM PFOS. Our verification of PFOS stock solution revealed contamination by PFNS, PFHpS, and PFHxS. As such, we assessed the body burden of these compounds in the PFOS-exposed larvae, as well. Statistics compared 48 hpf to 72 hpf of the same treatment group using unpaired t-tests. ± denotes standard deviation. n = 10 pooled whole larvae from 3 experimental replicates. \* $P$ <0.05; \*\*\* $P$ <0.001; \*\*\*\* $P$ <0.0001.

**Table S4.**
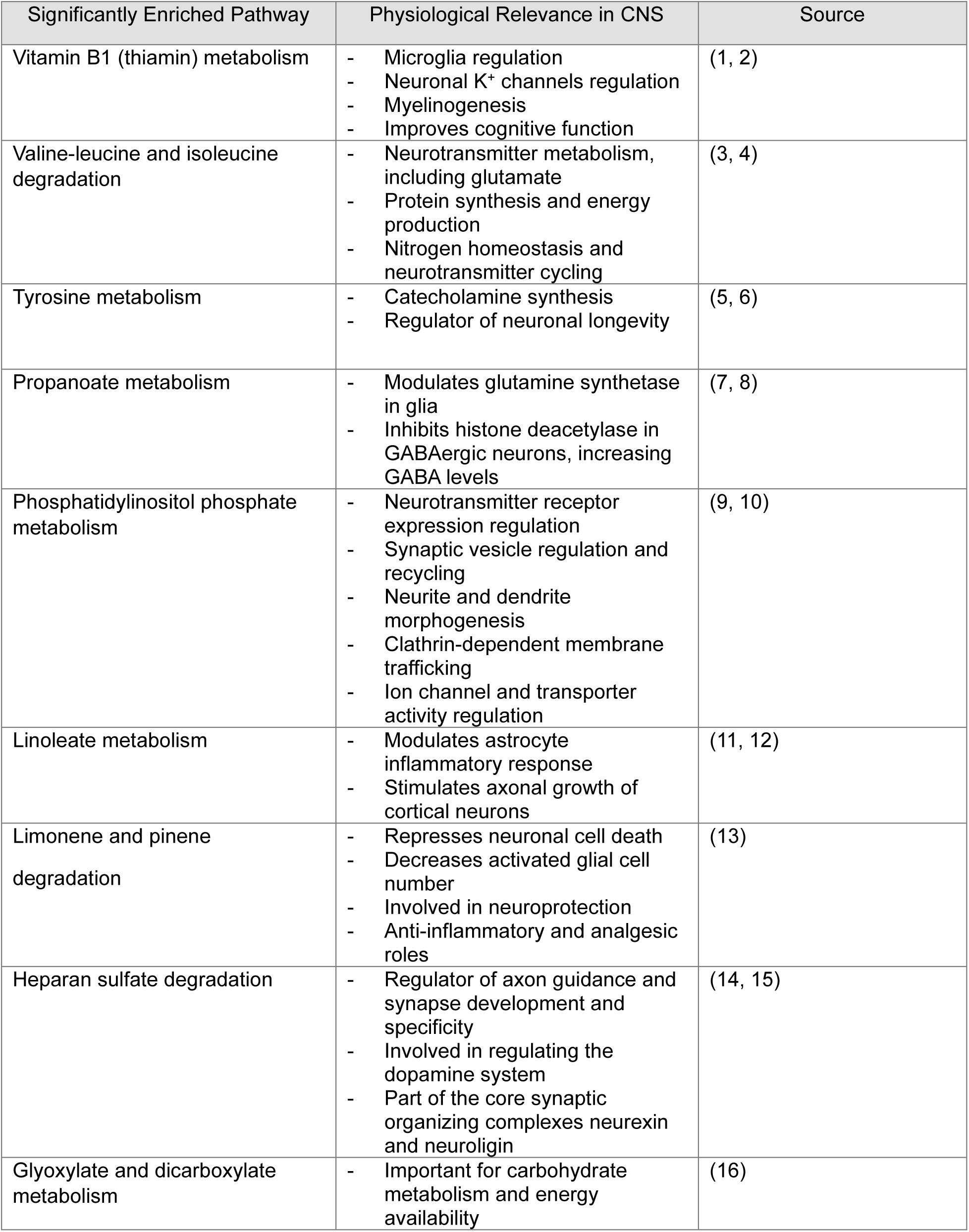
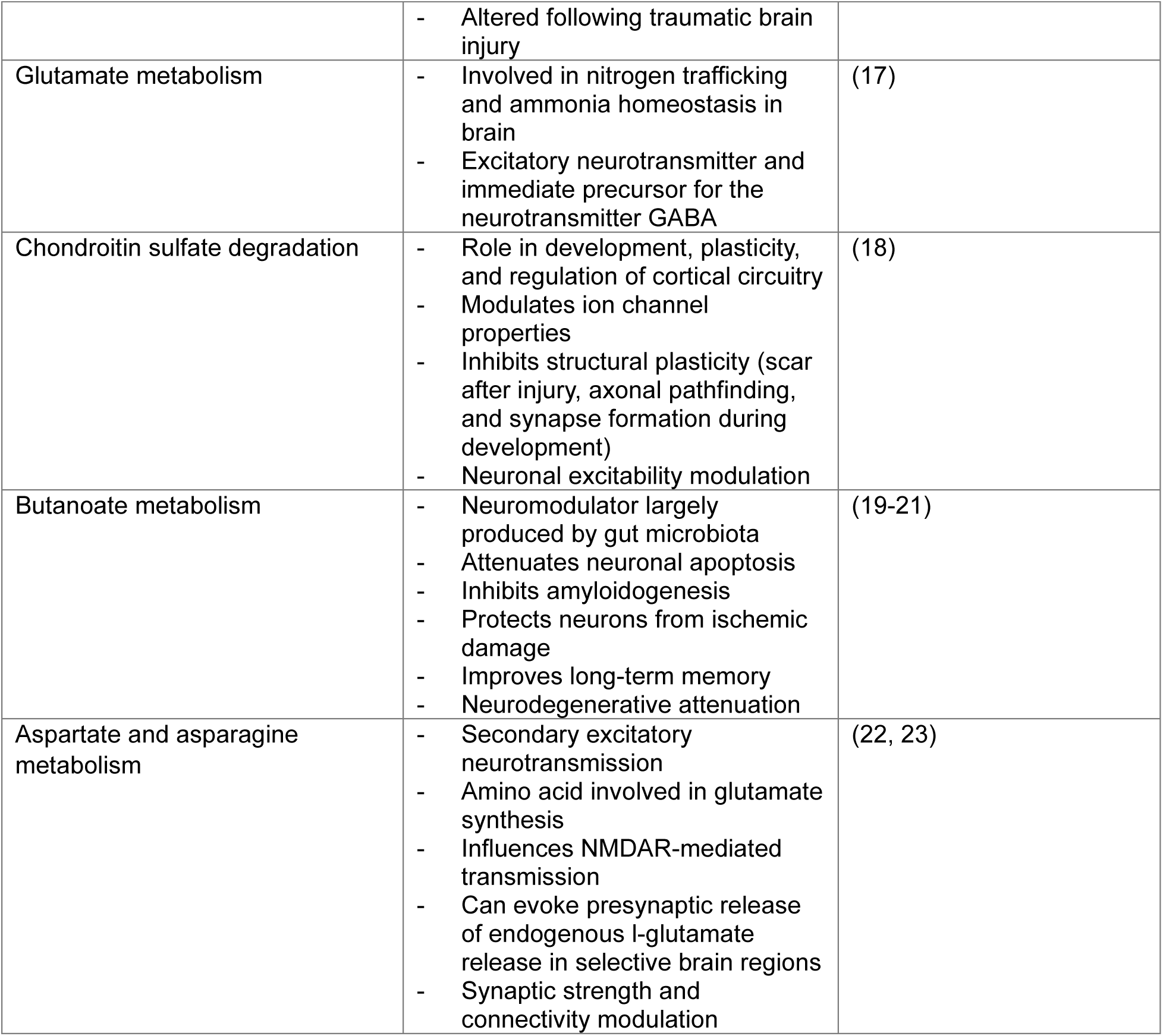
Significantly enriched metabolic pathways following 28 µM PFOS exposure in 3 dpf larvae.

### Supplemental Figures

**Figure S1.**
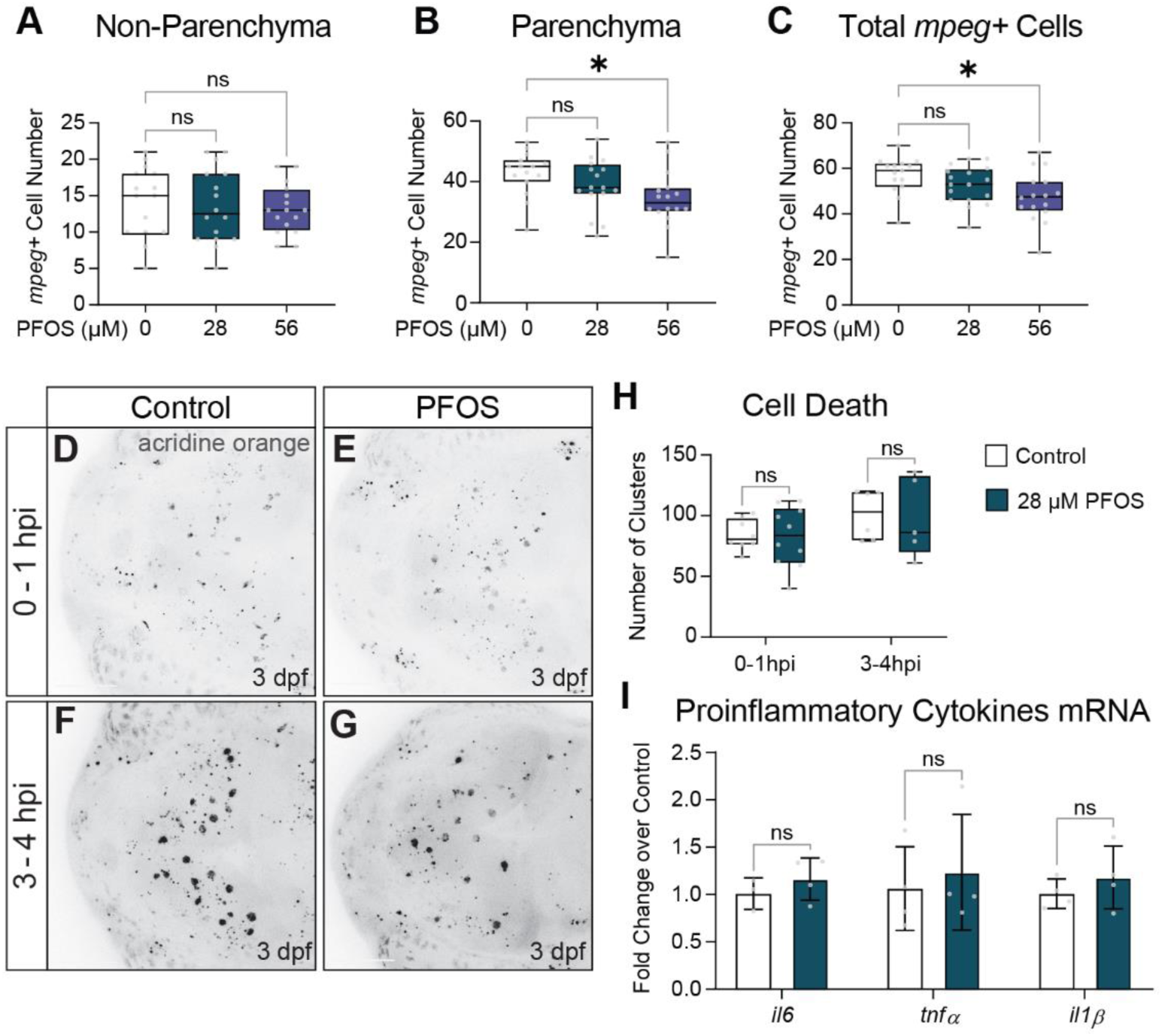
Microglia quantification, cell death, and proinflammatory cytokine mRNA analysis in PFOS-exposed larvae. (A) Non-parenchymal, (B) parenchymal, and (C) total microglia on were quantified in 3 dpf control (n = 15), 28 µM PFOS (n = 16), or 56 µM PFOS (n = 16) exposed larvae. There was no significant difference in parenchymal or non-parenchymal microglia number in 28 µM PFOS-exposed larvae compared to controls, while 56 µM PFOS larvae had significantly fewer parenchymal (*P* = 0.0168) and total microglia (*P* = 0.0205). (D,E) To assess cell death, live 3 dpf larvae were stained with 5 ug/mL acridine orange during the first hour post injury or (F,G) between 3-4 hpi. (H) Quantification of acridine orange-positive clusters in 3 dpf control and 28 µM PFOS-exposed larvae show no significant change in cell death (n = 5-9 per group). (I) qRT-PCR for the inflammatory genes *il6*, *tnfα*, and *il1β* were not significantly changed in isolated heads of 3 dpf control or 28 µM PFOS-exposed larvae (n = 10 pooled heads per sample). Confocal micrographs at 20x magnification. Box plot limits represent 25^th^ to 27^th^ percentile, with the midline representing the median. See Table S5 for additional statistical details.

**Figure S2.**
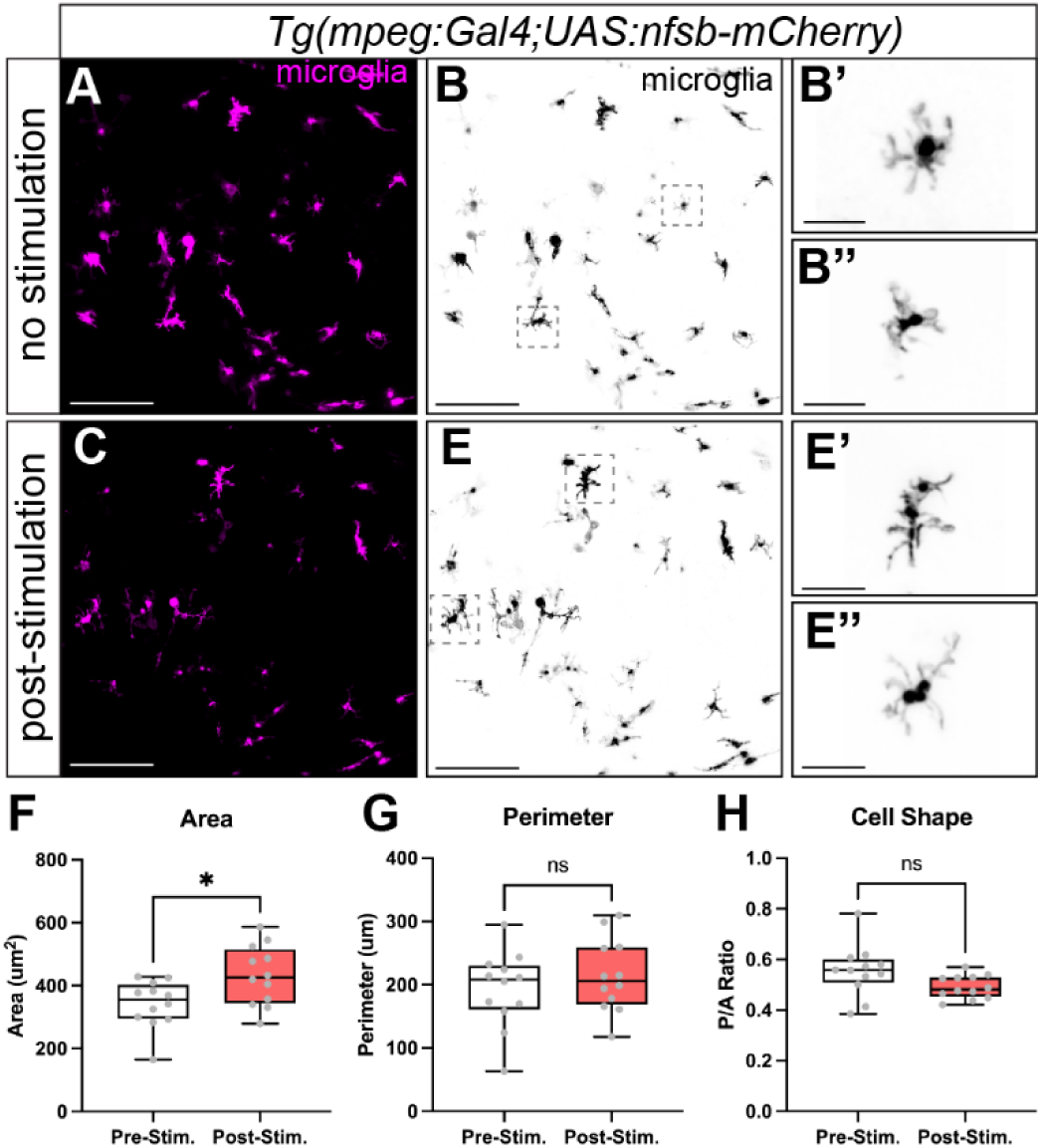
Control for the optogenetic microglia stimulation experiment by stimulating halorhodopsin negative microglia. To ensure that 589 nm light alone does not alter microglia morphology, PFOS-exposed transgenic larvae expressing macrophage-driven mCherry (*Tg(mpeg1:gal4FF;UAS:nfsb-mCherry)*) were either (A-B”) unstimulated or (C-E”) subjected to the 589 nm light for 4 hours. (F) While microglia area was increased following light exposure (*P* = 0.0168), (G) the light stimulation did not result in significant changes in perimeter or (H) the perimeter-to-area ratio. Therefore, light stimulation alone does not result in the ramification of microglia. n = 12 cells. Box plot limits represent 25^th^ to 27^th^ percentile, with the midline representing the median. See Table S5 for additional statistical details.

**Figure S3.**
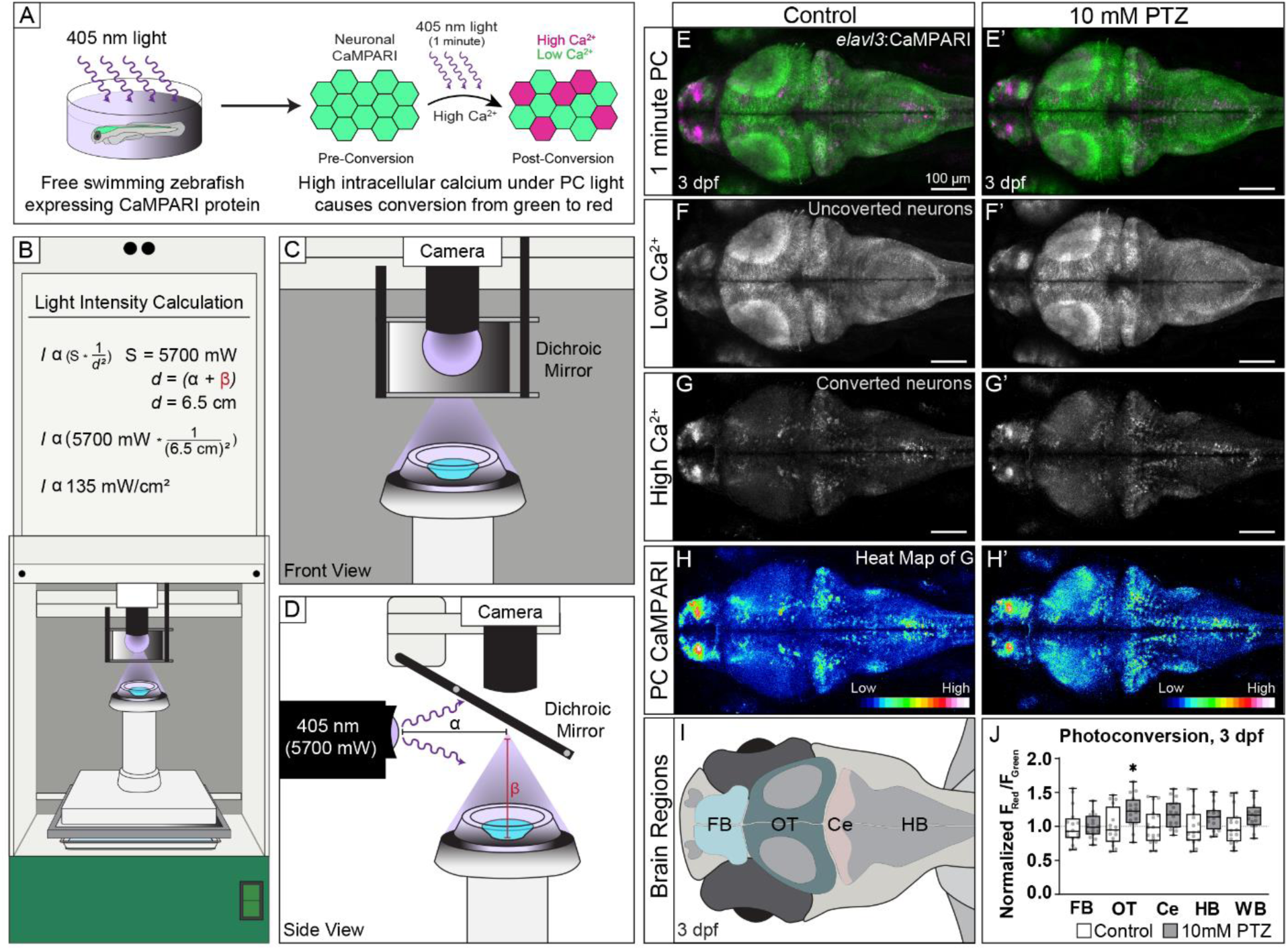
Functional neuroimaging to understand the effects of toxicant exposure on neuronal activity. (A) Schematic overview of CaMPARI photoconversion: free-swimming zebrafish larvae with neuron-specific expression of the genetically encoded calcium indicator CaMPARI protein (*Tg(elavl3:CaMPARI)*) are subjected to 405 nm light for 1 minute. Exposure to blue light photoconverts any neurons with high intracellular calcium from green to red. (B-D) We modified our Noldus DanioVision Behavioral unit, outfitted with LEDs for optogenetic manipulation, to efficiently photoconvert CaMPARI in free-swimming zebrafish. (B) A 3D printed pedestal is used to reduce the working distance between the light source and zebrafish larvae. Applying the inverse-square law, the light intensity at the apex of the pedestal was calculated to be 135 mW/cm². (C) Illustration of the apex of the pedestal containing a single well dish (OD 15 mm). (D) The 405 nm light has an intensity of 5700 mW at the source. The light travels horizontally then is reflected off a dichroic mirror within the unit. The working distance was calculated as the distance from the light source to the dichroic mirror (α) plus the distance from the dichroic mirror to the apex of the pedestal (β). (E) Confocal micrographs of neuronal calcium following 1-minute photoconversion of CaMPARI in 3 dpf larvae treated with egg water or (E’) after 10 minutes in 10 mM pentylenetetrazol (PTZ), a GABAA-inhibitor known to induce hyperactivity. (F, F’) Green, non-active neurons with low intracellular calcium from E & E’. (G, G’) Photoconverted, active neurons with high intracellular calcium, pseudo-labeled as magenta, from E & E’. (H, H’) An intensity LUTs applied to the active neurons in G & G’ spectrally maps the regional increases of neuronal activity on a scale from less active (blue) to more active (red-white). (I&J) Regional and global neuronal activity can be quantified by determining the ratio of fluorescent intensity in the red versus green channel (F_Red_/F_Green_). Exposure to 10 mM PTZ significantly increased brain activity in the optic tectum (OT; *P* = 0.0496) and trended toward increased activity in the cerebellum (Ce), hindbrain (HB), and whole brain (WB). Red/Green ratios were normalized relative to vehicle control. n = 16 fish per group. Box plot limits represent 25^th^ to 27^th^ percentile, with the midline representing the median. See Table S5 for additional statistical details.

**Figure S4.**
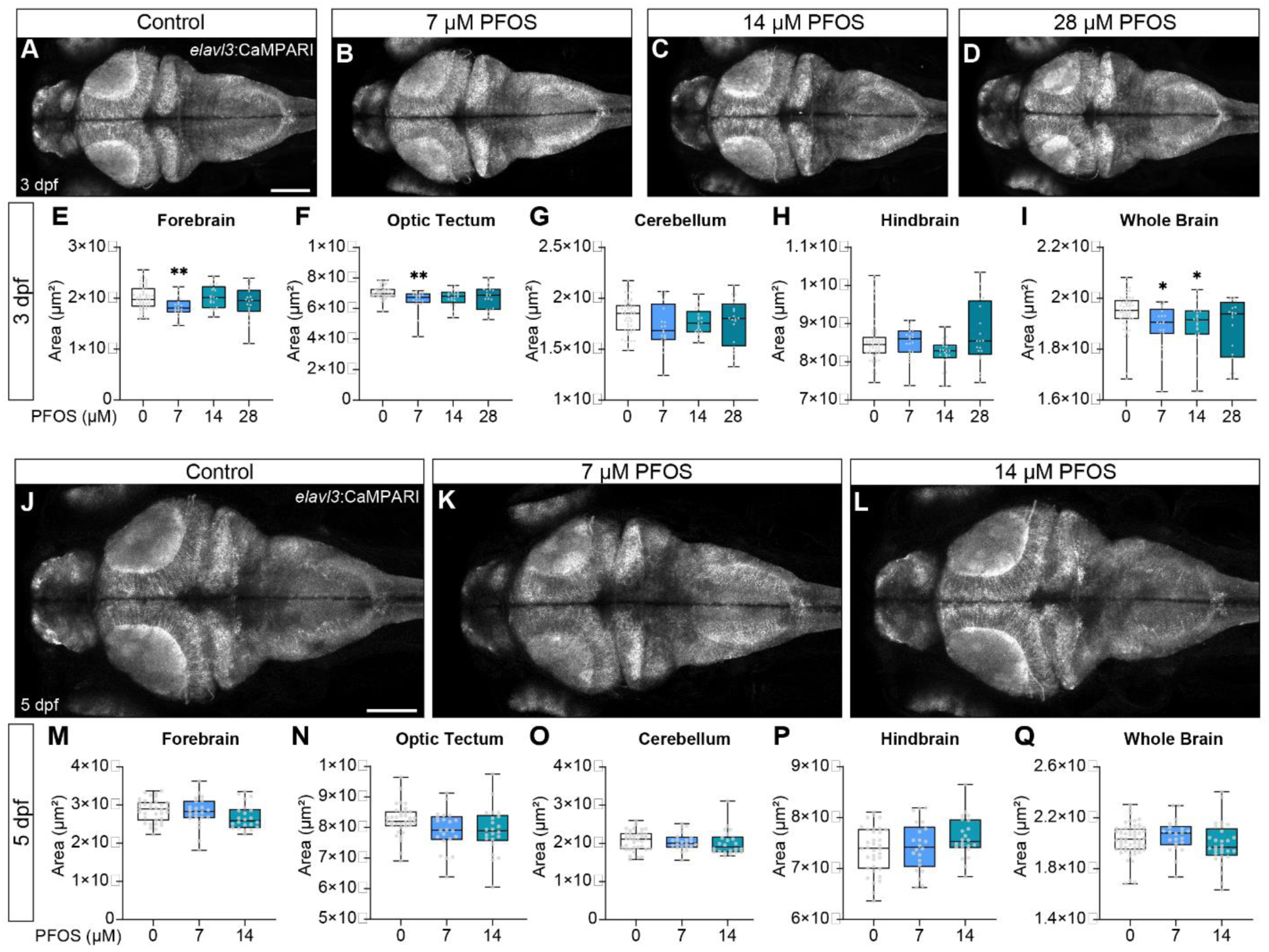
Regional and global brain morphology in larvae exposed to varying concentrations of PFOS. (A-D) Confocal micrographs of 3 dpf larvae or (J-L) 5 dpf larvae expressing *Tg(elavl3:CaMPARI*) exposed to (A&J) control, (B&K) 7 µM, (C&L) 14 µM, or (D) 28 µM PFOS. At 3 dpf, 7 µM PFOS larvae had a significant reduction in area of the (E) forebrain (Control vs 7 µM PFOS, *P =* 0.0035) and (F) optic tectum (Control vs 7 µM PFOS, *P =* 0.0047). (I) When measuring the area of the entire brain, only 7 µM and 14 µM PFOS exposure resulted in reduced global brain area (Control vs 7 µM & 14 µM, *P =* 0.0174 & *P =* 0.0373, respectively). At 5 dpf, there were no significant differences in regional area or global brain area with 7 µM nor 14 µM PFOS exposure. Confocal micrographs at 10x magnification. Control n = 52-54; treated n = 22-25. Scale bar = 100 um. Box plot limits represent 25^th^ to 27^th^ percentile, with the midline representing the median. See Table S5 for additional statistical details.

**Figure S5.**
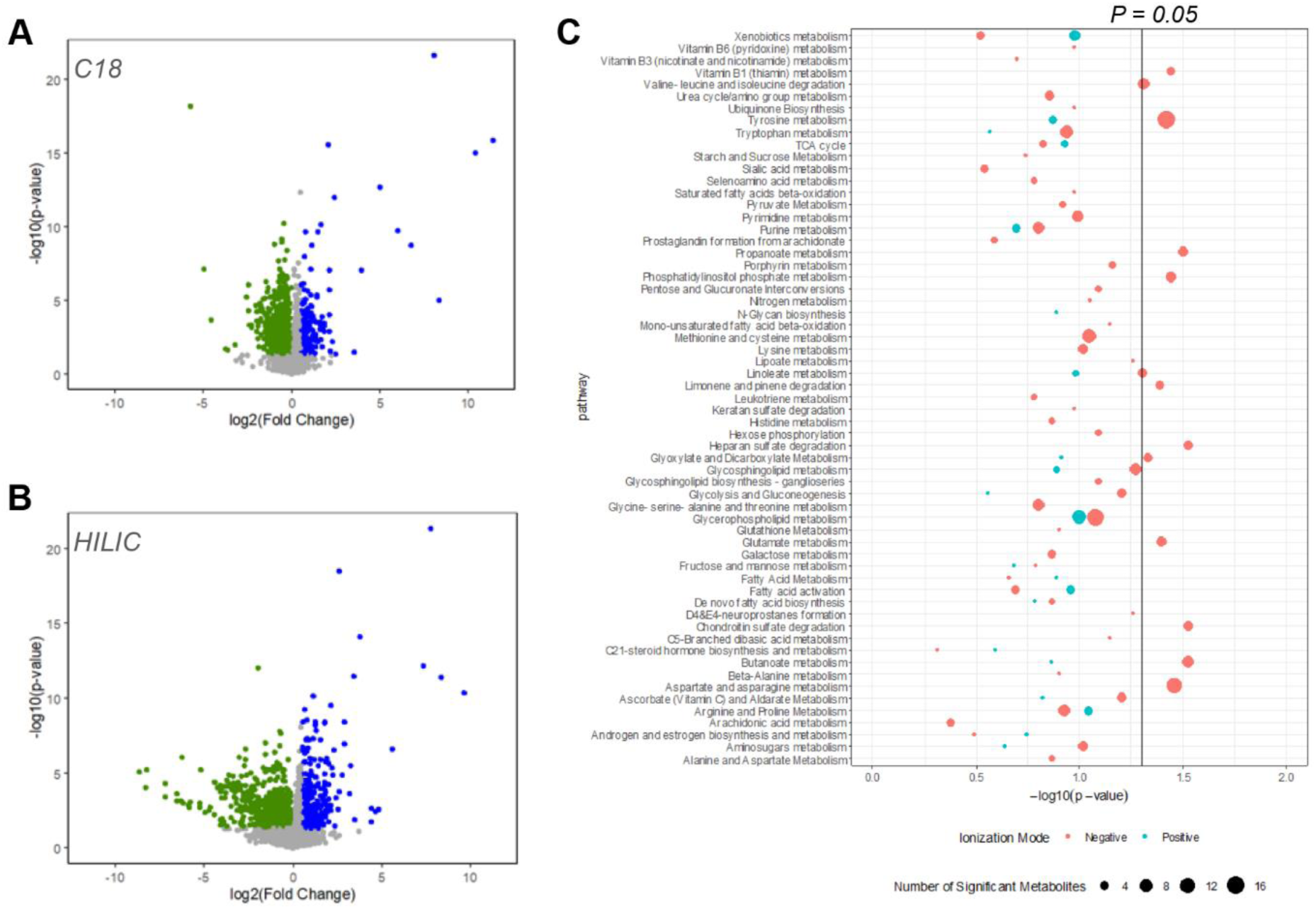
MWAS of heads collected from control versus PFOS-exposed larvae. Heads were collected from 3 dpf control or 28 µM PFOS exposed larvae for an untargeted metabolome wide association study (MWAS). (A) Volcano plots of metabolites detected using a C18 column with negative ionization and (B) a HILIC column with positive ionization reveal several significantly down- or up-regulated metabolites in the PFOS-exposed brain (Green = downregulated; Blue = upregulated). Statistical significance was set at *P* = 0.05. (C) There are several significantly enriched pathways following both negative ionization (pink) and positive ionization (teal). The vertical line is at *P* = 0.05. Size of each dot represents the number of significant metabolites in that pathway. HILIC, hydrophilic interaction liquid chromatography column.

**Figure S6.**
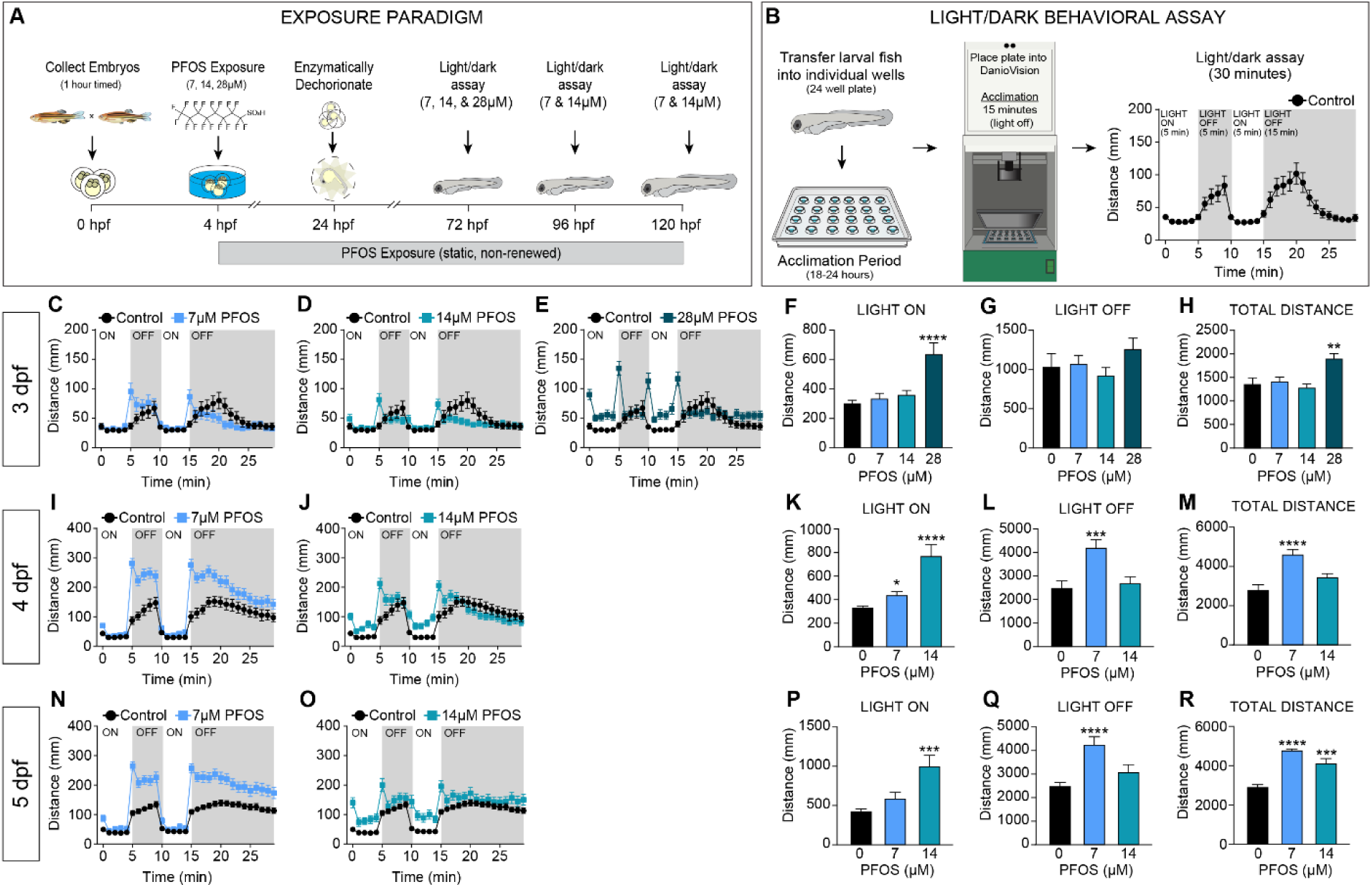
Light/dark behavioral assay in larvae exposed to varying concentrations of PFOS. (A) Exposure paradigm: at 4 hpf, zebrafish embryos were statically exposed to 0.1% DMSO (Control) or varying concentrations of PFOS (7, 14, or 28 µM). At 24 hpf, embryos were enzymatically dechorionated. Behavior was captured at 72, 96, and 120 hpf. (B) Light/dark behavioral assay: Zebrafish larvae were placed in individual wells of a 24-well plate containing the relevant toxicant solution 18-24 hours prior to the behavior assay to allow for well acclimation. A light/dark behavioral assay was performed using the Noldus DanioVision Behavioral Unit. The light/dark behavioral assay was as follows: 15-minute acclimation in the unit with the light off, 5 minutes with the light on, 5 minutes with the light off, 5 minutes with the light on, and 15 minutes with the light off. (C-H) At 3 dpf, larvae exposed to 28 µM PFOS had a significant increase in swim activity during the (F) light on cycles (*P* = 0.0003) and (H) over the course of the assay (*P* = 0.0123). (K) At 4 dpf, 7 µM PFOS larvae had a significant increase in swim distance when the light was on (*P* = 0.0277), as did 14 µM PFOS larvae (*P* = 0.0002). (L) At 4 dpf, 7 µM PFOS-exposed larvae also had a significant increase in swim behavior when the light was off (*P* = 0.0018) and (M) in total over the course of the assay (*P* < 0.0001). (P) At 5dpf, 14 µM PFOS had increased swim behavior when the light was on (*P* = 0.0006), and (Q) 7 µM PFOS larvae had increased swim behavior when the light was off (*P* < 0.0001). (R) Both 7 µM (*P* < 0.0001) and 14 µM PFOS (*P* = 0.0003) larvae had a significant increase in total distance moved. Control n = 130-190, treated n = 88-99. Error bars represent SEM. See Table S5 for additional statistical details.

**Figure S7.**
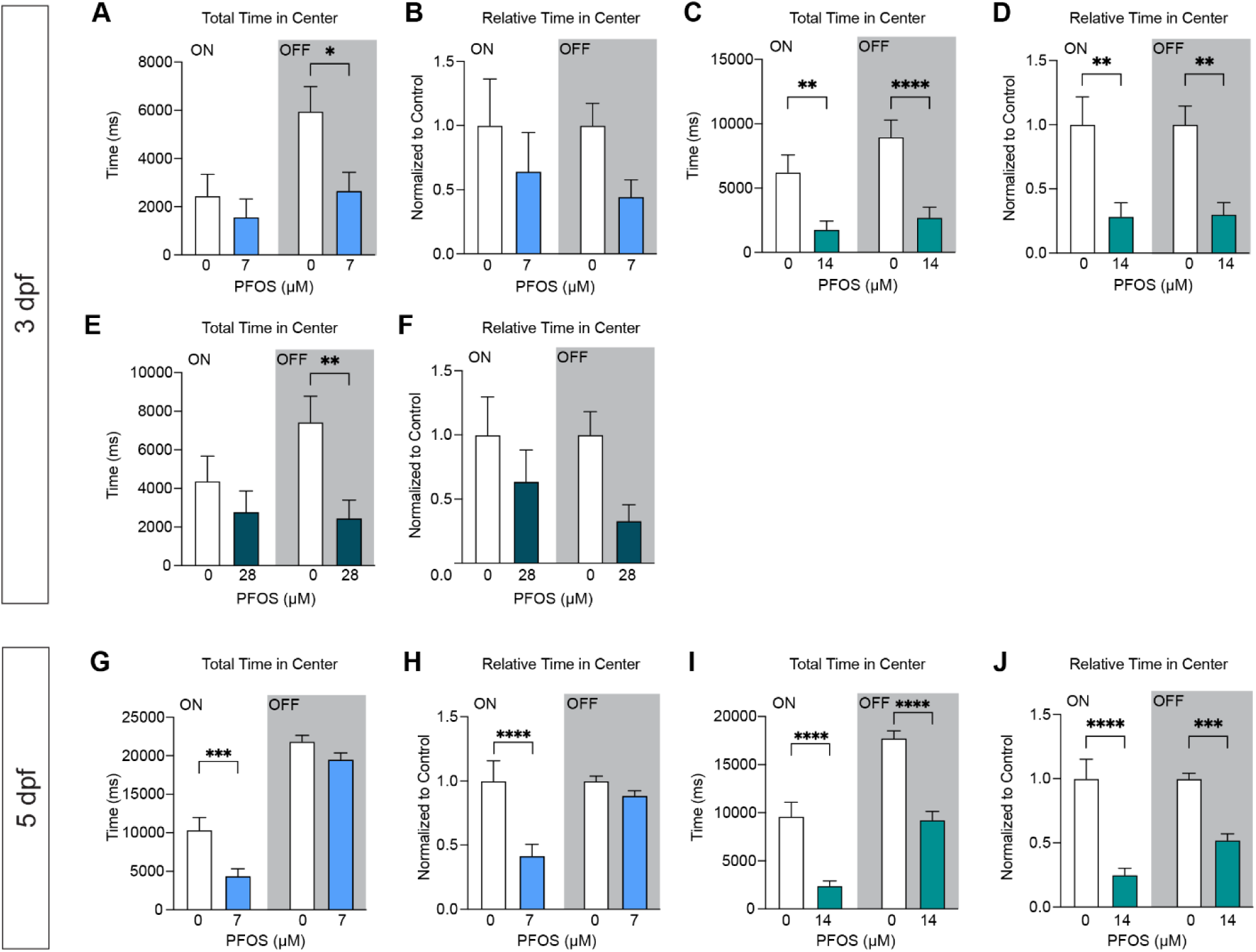
Absolute and relative time spent in center throughout the light/dark behavioral response assay. (A) Total time spent in the center during the light on and light off cycles at 3dpf, control versus 7 µM (*P* = 0.0139, “OFF”). (B) Relative time spent in the center during the light on and light off cycles at 3 dpf, control versus 7 µM. (C) Total time spent in the center during the light on and light off cycles at 3 dpf, control versus 14 µM (*P* = 0.0075, “ON”; *P* < 0.0001, “OFF”). (D) Relative time spent in the center during the light on and light off cycles at 3 dpf, control versus 14 µM (*P* = 0.0015, “ON”; *P* = 0.0018, “OFF”). (E) Total time spent in the center during the light on and light off cycles at 3 dpf, control versus 28 µM (*P* = 0.0061, “OFF”). (F) Relative time spent in the center during the light on and light off cycles at 3 dpf, control versus 28 µM. (G) Total time spent in the center during the light on and light off cycles at 5 dpf, control versus 7 µM (*P* = 0.0004, “ON”). (H) Relative time spent in the center during the light on and light off cycles at 5 dpf, control versus 7 µM (*P* < 0.0001). (I) Total time spent in the center during the light on and light off cycles at 5 dpf, control versus 14 µM (*P* < 0.0001, “ON”; *P* < 0.0001, “OFF”). (J) Relative time spent in the center during the light on and light off cycles at 5 dpf, control versus 14 µM (*P* < 0.0001, “ON”; *P* = 0.0004, “OFF”). Control n = 88-102 for controls; treated n = 84-100. Error bars represent SEM. See Table S5 for additional statistical details.

**Figure S8.**
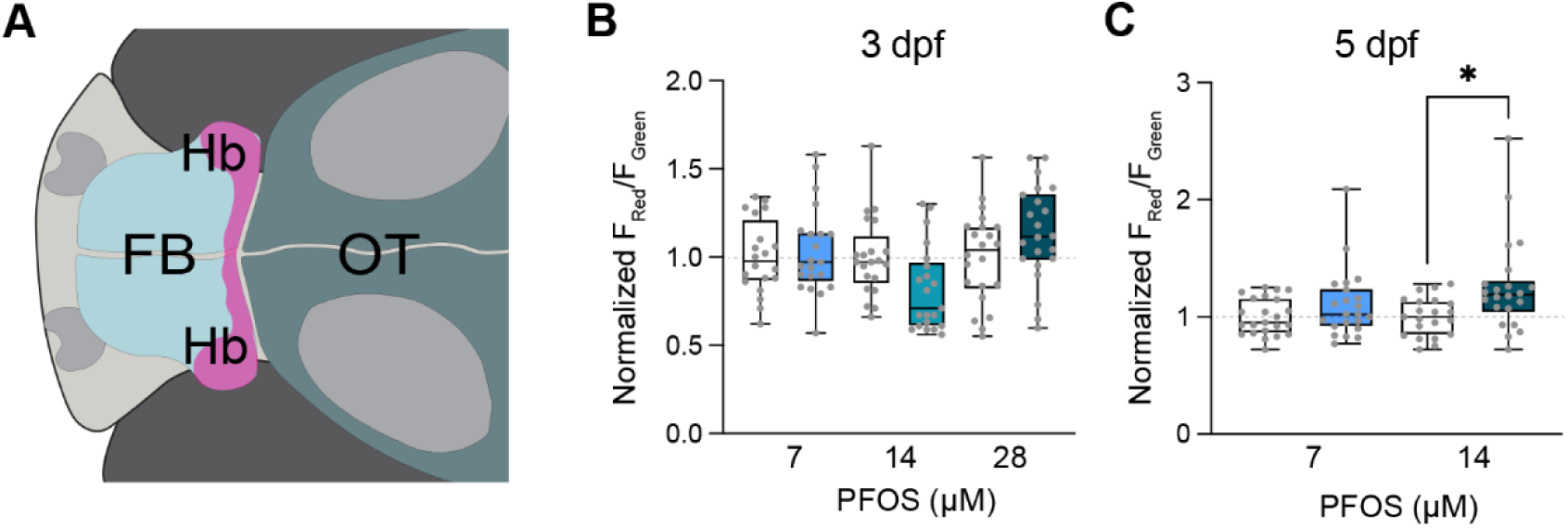
CaMPARI analysis of the habenula in larvae exposed to varying concentrations to PFOS. (A) Illustrative representation of a larval zebrafish brain with anatomical regions outlined: forebrain (FB), habenula (Hb), and the optic tectum (OT). Activity quantification of neuron-driven CaMPARI was performed in the entire developing habenula following 1-minute exposure to 405 nm light. (B) At 3 dpf, there were no significant differences in habenular Ca2+ activity. (C) At 5 dpf, 14 µM PFOS-exposed larvae had a significant increase in habenular Ca^2+^ activity (*P* = 0.0170), but 7 µM PFOS was unchanged. n = 20-22 fish per treatment. Box plot limits represent 25^th^ to 27^th^ percentile, with the midline representing the median. See Table S5 for additional statistical details.

**Figure S9.**
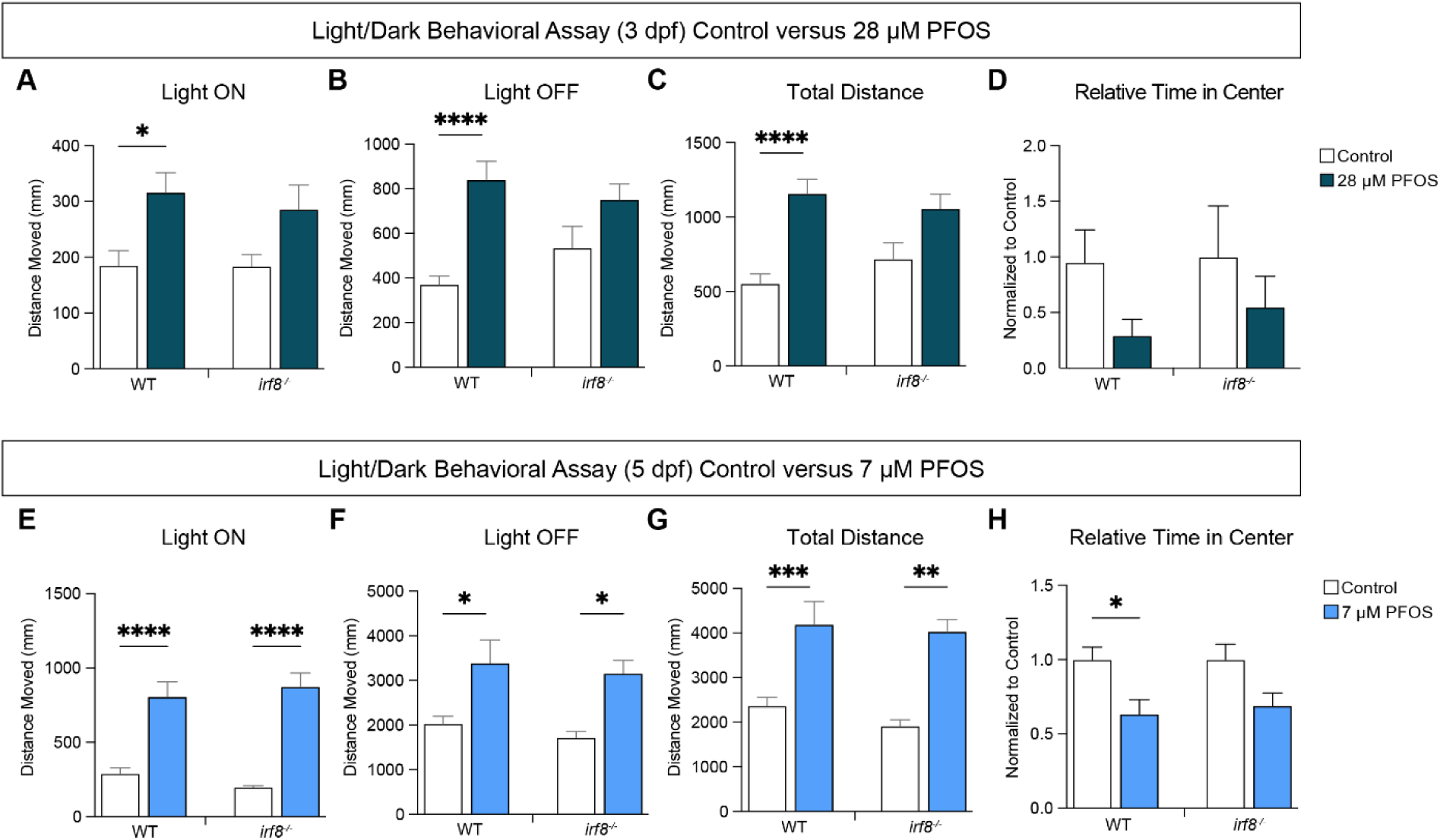
Distance traveled in control versus PFOS-treated wildtype and macrophage mutant larvae. (A) 3 dpf light on (*P* = 0.0297, WT; *P* = 0.3036, *irf8^-/-^*). (B) 3dpf light off (*P* < 0.0001, WT; *P* = 0.2162, *irf8^-/-^*). (C) 3 dpf total distance moved throughout assay (*P* < 0.0001, WT; *P* = 0.1069, *irf8^-/-^*). (D) Relative time spent in center normalized to the respective control groups. (E) 5 dpf light on (*P* < 0.0001, WT; *P* < 0.0001, *irf8^-/-^*). (F) 5 dpf light off (*P* = 0.0200, WT; *P* = 0.0456, *irf8^-/-^*). (G) 5 dpf total distance moved throughout assay (*P* = 0.0009, WT; *P* = 0.0011, *irf8^-/-^*). (H) Relative time spent in center normalized to the respective control groups (*P* = 0.0213, WT; *P* = 0.1513, *irf8^-/-^*). n = 14-22 per group. Error bars represent SEM. See Table S5 for additional statistical details.

**Figure S10.**
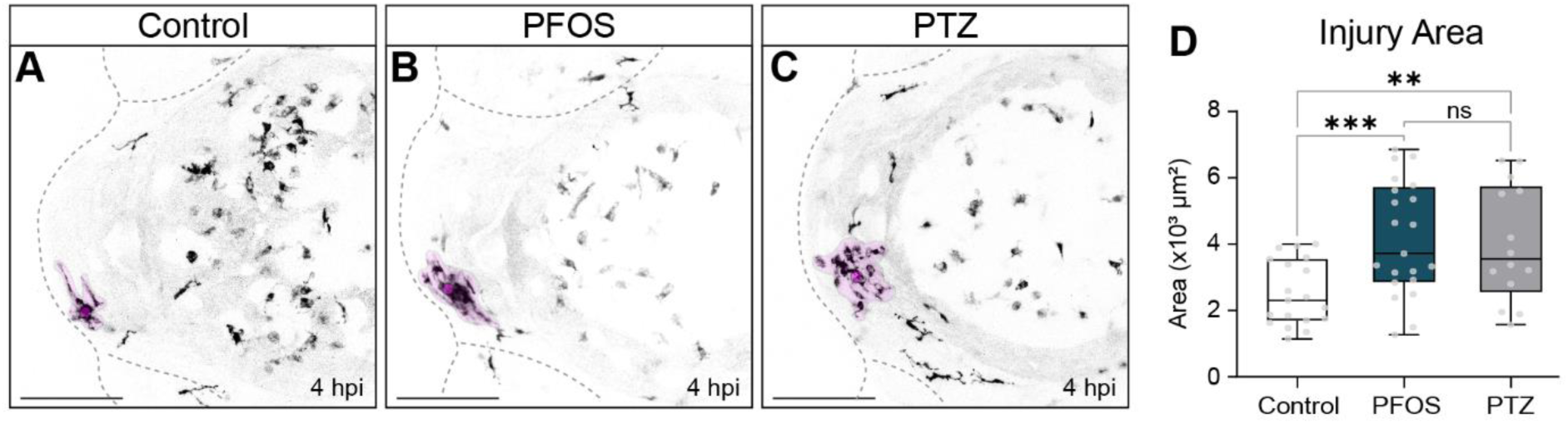
Microglia response to brain injury in control versus PFOS-exposed versus PTZ-exposed larvae at 3 dpf. Larvae with transgenic expression of macrophages (Tg(*mpeg1:EGFP*)) were dosed with either (A) control or (B) 28 µM PFOS at 4 hpf, or (C) 5 mM PTZ at 72 hpf, followed by brain injury. (D) At 4 hpi, PFOS-exposed larvae had a significant increase in microglia response area (*P* = 0.0082). In addition, neuronal excitation by PTZ also resulted in a significant increase in microglia response (*P* = 0.0314). n = 14-21 per group. Box plot limits represent 25^th^ to 27^th^ percentile, with the midline representing the median. See Table S5 for additional statistical details.

**Figure S11.**
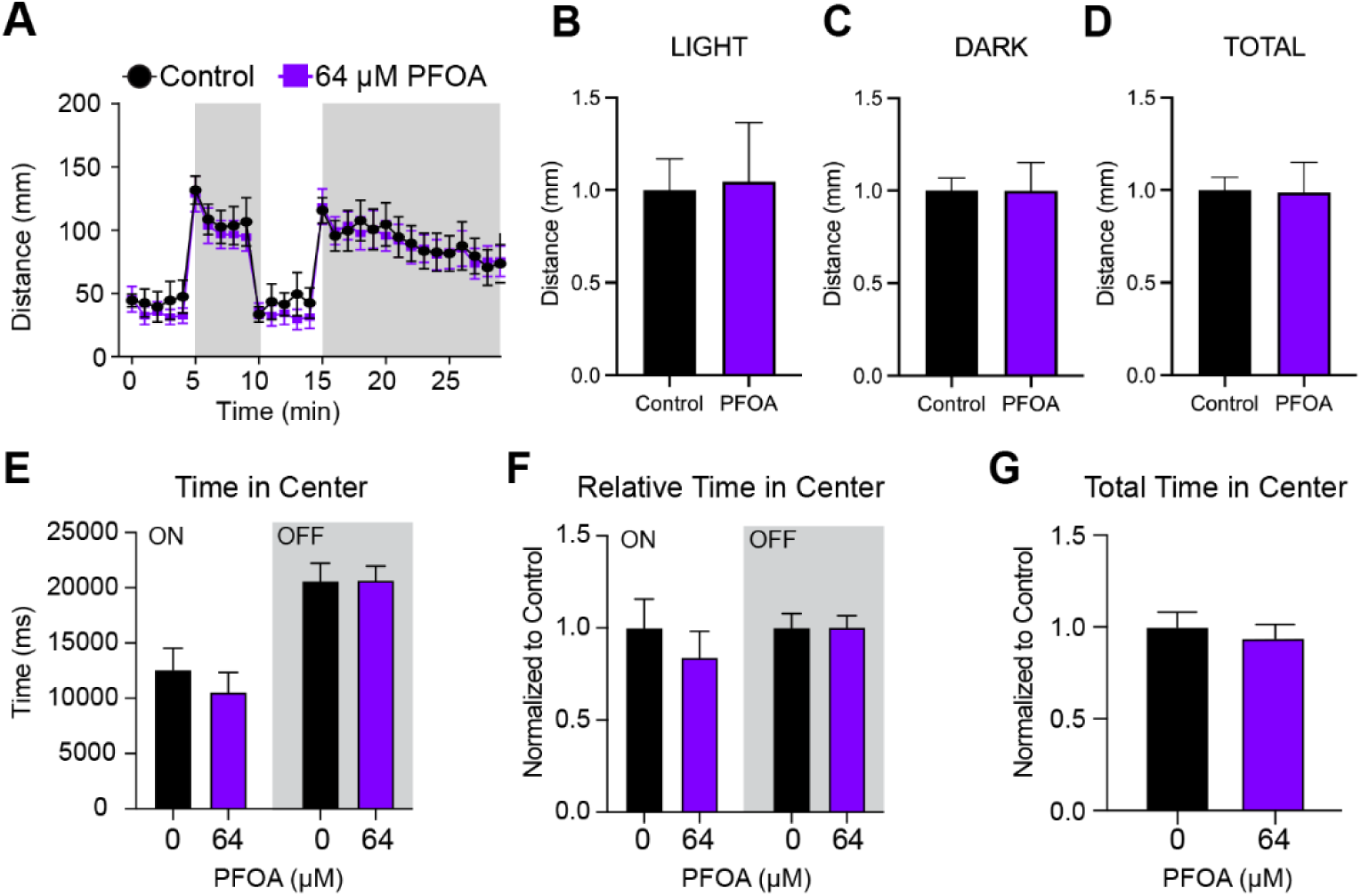
Light/dark behavioral analysis in PFOA-exposed larvae. Zebrafish larvae were subjected to a 30-minute light/dark behavioral assay at 5 dpf, as previously described. (A) There were no apparent changes in distance moved during the 30-minute assay. (B) Total distance moved during light cycles, (C) dark cycles, and (D) in total throughout the assay were not significantly changed. (E-G) PFOA exposure also did not result in changes in center-avoidance behavior during the light on or light off cycles. n = 24 control and n = 46 treated for behavior analyses; n = 37 control and n = 35 treated for anxiety analyses. See Table S5 for additional statistical details.

**Figure S12.**
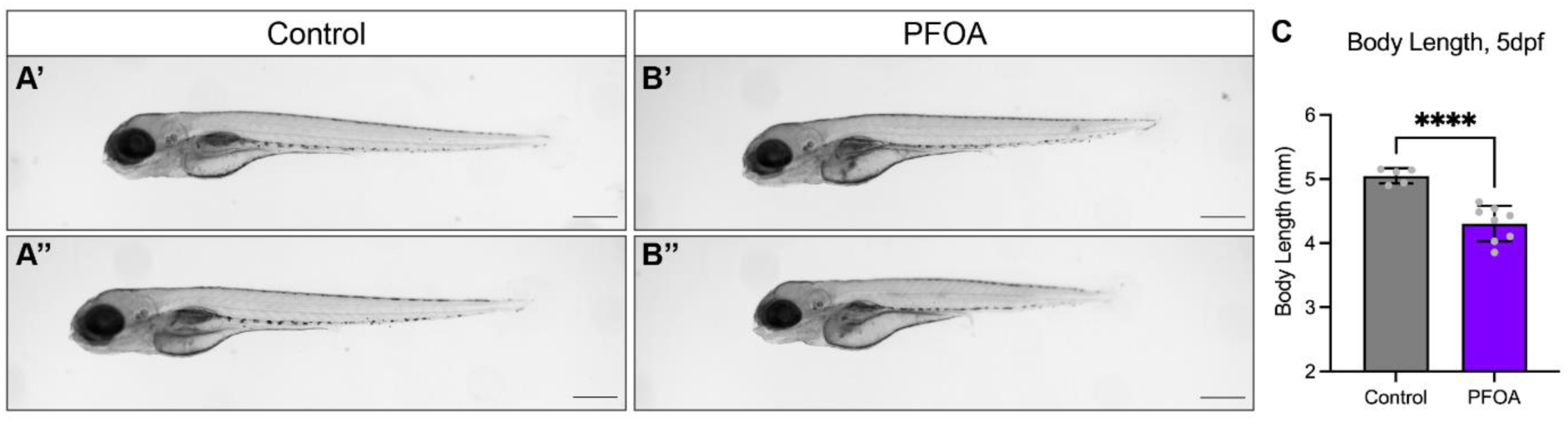
Body length of 5 dpf exposed to PFOA. (A’-A”) Images of 5 dpf control-treated larvae. (B’- B”) Images of 5 dpf larvae chronically treated with 64 µM PFOA. (C) Body length measurements of control versus PFOA treatment at 5 dpf (*P* < 0.0001). n = 5-8 per group. Scale = 500 um. See Table S5 for additional statistical details.

