## Supplemental Table S5 for "Evaluation of neural regulation and microglial responses to brain injury in larval zebrafish exposed to perfluorooctane sulfonate"

**Table S5. Detailed statistical information for each figure**

| **Figure** | **Statistical Test** | **P Value** | **Sample size (n)** | **Age** | **Treatment** | **Transgenic line** |
| --- | --- | --- | --- | --- | --- | --- |
| Figure 1N | Unpaired parametric t-test with Welch's Correction | 0.0342 | 15 fish, 3-18 cells counted per fish | 3 dpf | 0.1% DMSO; 28 µM PFOS | *Tg(HuC:Kaede; mpeg1:EGFP)*  *Some larvae also used* |
| FIgure 1O | Unpaired parametric t-test with Welch's Correction | 0.0006 | 15 fish, 3-18 cells counted per fish | 3 dpf | 0.1% DMSO; 28 µM PFOS | *Tg(HuC:Kaede; mpeg1:EGFP)* |
| Figure 1P | Unpaired parametric t-test with Welch's Correction | 0.0002 | 15 fish, 3-18 cells counted per fish | 3 dpf | 0.1% DMSO; 28 µM PFOS | *Tg(HuC:Kaede; mpeg1:EGFP)* |
| Figure 1Q | Unpaired parametric t-test with Welch's Correction | 0.0211 | 4 replicates of 10 pooled heads | 3 dpf | 0.1% DMSO; 28 µM PFOS | *Tg(mpeg1:EGFP)* |
| Figure 1R | Unpaired parametric t-test with Welch's Correction | 0.0208 | 5-6 per group | 3 dpf | 0.1% DMSO; 28 µM PFOS | *Tg(mpeg1:EGFP)* |
| Figure 1S | Unpaired non-parametric t-test (Mann-Whitney Wilcoxon) | 0.0037 | 19-20 per group | 3 dpf | 0.1% DMSO; 28 µM PFOS | *Tg(mpeg1:EGFP)* |
| Figure 1T | Unpaired parametric t-test with Welch's Correction | 0.0467 | 4 per group | 3 dpf | 0.1% DMSO; 28 µM PFOS | *Tg(mpeg1:EGFP)* |
| Figure S1A | Non-parametric ANOVA (Kruskal-Wallis test) with Dunn's test for multiple comparisons | see figure caption | 15-16 per group | 3 dpf | 0.1% DMSO; 28 µM PFOS; 56 µM PFOS | *Tg(HuC:Kaede; mpeg1:EGFP)* |
| Figure S1B | Welch ANOVA with post-hoc Dunnett's T3 multiple comparisons test | see figure caption | 15-16 per group | 3 dpf | 0.1% DMSO; 28 µM PFOS; 56 µM PFOS | *Tg(HuC:Kaede; mpeg1:EGFP)* |
| Figure S1C | Welch ANOVA with post-hoc Dunnett's T3 multiple comparisons test | see figure caption | 15-16 per group | 3 dpf | 0.1% DMSO; 28 µM PFOS; 56 µM PFOS | *Tg(HuC:Kaede; mpeg1:EGFP)* |
| Figure S1H | Two-Way ANOVA with post-hoc Tukey multiple comparisons test | see figure caption | 5-9 per group | 3 dpf | 0.1% DMSO; 28 µM PFOS | *Tg(mpeg1:EGFP)* |
| Figure S1I | Multiple unpaired parametric t-tests with post-hoc FDR analysis for multple comparisons | see figure caption | 4 replicates of 10 pooled heads | 3 dpf | 0.1% DMSO; 28 µM PFOS | *Tg(mpeg1:EGFP)* |
| Figure 2L | Unpaired non-parametric t-test (Mann-Whitney Wilcoxon) | 0.5316 | 40-42 cells per group across 3 experimental replicates | 3 dpf | 28 µM PFOS | *Tg(mpeg1:GalFF;*  *UAS:eNpHR3.0-mCherry)* |
| Figure 2M | Unpaired non-parametric t-test (Mann-Whitney Wilcoxon) | 0.007 | 40-42 cells per group across 3 experimental replicates | 3 dpf | 28 µM PFOS | *Tg(mpeg1:GalFF;*  *UAS:eNpHR3.0-mCherry)* |
| Figure 2N | Unpaired non-parametric t-test (Mann-Whitney Wilcoxon) | <0.0001 | 40-42 cells per group across 3 experimental replicates | 3 dpf | 28 µM PFOS | *Tg(mpeg1:GalFF;*  *UAS:eNpHR3.0-mCherry)* |
| Figure 2O | Two-Way ANOVA with post-hoc Sidak multiple comparisons test | see figure | 9-15 per group | 3 dpf | 0.1% DMSO; 28 µM PFOS | *Tg(mpeg1:GalFF;*  *UAS:eNpHR3.0-mCherry)* |
| Figure S2F | Unpaired parametric t-test with Welch's Correction | 0.0168 | 12 cells per group across 3 experimental replicates | 3 dpf | 28 µM PFOS | *Tg(mpeg1:gal4FF;*  *UAS:nfsb-mCherry)* |
| Figure S2G | Unpaired parametric t-test with Welch's Correction | 0.3841 | 12 cells per group across 3 experimental replicates | 3 dpf | 28 µM PFOS | *Tg(mpeg1:gal4FF;*  *UAS:nfsb-mCherry)* |
| Figure S2H | Unpaired parametric t-test with Welch's Correction | 0.0686 | 12 cells per group across 3 experimental replicates | 3 dpf | 28 µM PFOS | *Tg(mpeg1:gal4FF;*  *UAS:nfsb-mCherry)* |
| Figure 3H | Non-parametric ANOVA (Kruskal-Wallis test) with Dunn's test for multiple comparisons | see figure | 21-23 per group | 3 dpf | 0.1% DMSO; 7 µM PFOS | *Tg(elavl1:CaMPARI)* |
| Figure 3I | Non-parametric ANOVA (Kruskal-Wallis test) with Dunn's test for multiple comparisons | see figure | 21-23 per group | 3 dpf | 0.1% DMSO; 28 µM PFOS | *Tg(elavl1:CaMPARI)* |
| Figure 3M | Non-parametric ANOVA (Kruskal-Wallis test) with Dunn's test for multiple comparisons | see figure | 21-23 per group | 5 dpf | 0.1% DMSO; 7 µM PFOS | *Tg(elavl1:CaMPARI)* |
| Figure 3N | Non-parametric ANOVA (Kruskal-Wallis test) with Dunn's test for multiple comparisons | see figure | 21-23 per group | 5 dpf | 0.1% DMSO; 14 µM PFOS | *Tg(elavl1:CaMPARI)* |
| Figure S3J | Non-parametric ANOVA (Kruskal-Wallis test) with Dunn's test for multiple comparisons | see figure | 16 per group | 3 dpf | 10 mM PTZ | *Tg(elavl1:CaMPARI)* |
| Figure S4E | Welch ANOVA with post-hoc Dunnett's T3 multiple comparisons test | see figure | Control = 52-54; PFOS = 22-25 | 3 dpf | 0.1% DMSO; 7 µM PFOS; 14 µM PFOS; 28 µM PFOS | *Tg(elavl3:CaMPARI*) |
| Figure S4F | Non-parametric ANOVA (Kruskal-Wallis test) with Dunn's test for multiple comparisons | see figure | Control = 52-54; PFOS = 22-25 | 3 dpf | 0.1% DMSO; 7 µM PFOS; 14 µM PFOS; 28 µM PFOS | *Tg(elavl3:CaMPARI*) |
| Figure S4G | Welch ANOVA with post-hoc Dunnett's T3 multiple comparisons test | see figure | Control = 52-54; PFOS = 22-25 | 3 dpf | 0.1% DMSO; 7 µM PFOS; 14 µM PFOS; 28 µM PFOS | *Tg(elavl3:CaMPARI*) |
| Figure S4H | Non-parametric ANOVA (Kruskal-Wallis test) with Dunn's test for multiple comparisons | see figure | Control = 52-54; PFOS = 22-25 | 3 dpf | 0.1% DMSO; 7 µM PFOS; 14 µM PFOS; 28 µM PFOS | *Tg(elavl3:CaMPARI*) |
| Figure S4I | Non-parametric ANOVA (Kruskal-Wallis test) with Dunn's test for multiple comparisons | see figure | Control = 52-54; PFOS = 22-25 | 3 dpf | 0.1% DMSO; 7 µM PFOS; 14 µM PFOS; 28 µM PFOS | *Tg(elavl3:CaMPARI*) |
| Figure S4M | Welch ANOVA with post-hoc Dunnett's T3 multiple comparisons test | see figure | Control = 52-54; PFOS = 22-25 | 5 dpf | 0.1% DMSO; 7 µM PFOS; 14 µM PFOS | *Tg(elavl3:CaMPARI*) |
| Figure S4N | Non-parametric ANOVA (Kruskal-Wallis test) with Dunn's test for multiple comparisons | see figure | Control = 52-54; PFOS = 22-25 | 5 dpf | 0.1% DMSO; 7 µM PFOS; 14 µM PFOS | *Tg(elavl3:CaMPARI*) |
| Figure S4O | Non-parametric ANOVA (Kruskal-Wallis test) with Dunn's test for multiple comparisons | see figure | Control = 52-54; PFOS = 22-25 | 5 dpf | 0.1% DMSO; 7 µM PFOS; 14 µM PFOS | *Tg(elavl3:CaMPARI*) |
| Figure S4P | Welch ANOVA with post-hoc Dunnett's T3 multiple comparisons test | see figure | Control = 52-54; PFOS = 22-25 | 5 dpf | 0.1% DMSO; 7 µM PFOS; 14 µM PFOS | *Tg(elavl3:CaMPARI*) |
| Figure S4Q | Non-parametric ANOVA (Kruskal-Wallis test) with Dunn's test for multiple comparisons | see figure | Control = 52-54; PFOS = 22-25 | 5 dpf | 0.1% DMSO; 7 µM PFOS; 14 µM PFOS | *Tg(elavl3:CaMPARI*) |
| Figure 4C | Welch's ANOVA with post-hoc Games-Howell's multiple comparisons test | see figure | 76-101 per group | 3 dpf | 0.1% DMSO; 7 µM PFOS; 14 µM PFOS; 28 µM PFOS | AB |
| Figure 4D | Non-parametric ANOVA (Kruskal-Wallis test) with Dunn's test for multiple comparisons | see figure | 76-101 per group | 5 dpf | 0.1% DMSO; 7 µM PFOS; 14 µM PFOS | AB |
| Figure S6F | Welch's ANOVA with post-hoc Games-Howell's multiple comparisons test | see figure | Control = 130-190; PFOS = 88-99 | 3 dpf | 0.1% DMSO; 7 µM PFOS; 14 µM PFOS; 28 µM PFOS | AB |
| Figure S6G | Welch's ANOVA with post-hoc Games-Howell's multiple comparisons test | see figure | Control = 130-190; PFOS = 88-99 | 3 dpf | 0.1% DMSO; 7 µM PFOS; 14 µM PFOS; 28 µM PFOS | AB |
| Figure S6H | Welch's ANOVA with post-hoc Games-Howell's multiple comparisons test | see figure | Control = 130-190; PFOS = 88-99 | 3 dpf | 0.1% DMSO; 7 µM PFOS; 14 µM PFOS; 28 µM PFOS | AB |
| Figure S6K | Welch's ANOVA with post-hoc Games-Howell's multiple comparisons test | see figure | Control = 130-190; PFOS = 88-99 | 4 dpf | 0.1% DMSO; 7 µM PFOS; 14 µM PFOS | AB |
| Figure S6L | Welch's ANOVA with post-hoc Games-Howell's multiple comparisons test | see figure | Control = 130-190; PFOS = 88-99 | 4 dpf | 0.1% DMSO; 7 µM PFOS; 14 µM PFOS | AB |
| Figure S6M | Welch's ANOVA with post-hoc Games-Howell's multiple comparisons test | see figure | Control = 130-190; PFOS = 88-99 | 4 dpf | 0.1% DMSO; 7 µM PFOS; 14 µM PFOS | AB |
| Figure S6P | Welch's ANOVA with post-hoc Games-Howell's multiple comparisons test | see figure | Control = 130-190; PFOS = 88-99 | 5 dpf | 0.1% DMSO; 7 µM PFOS; 14 µM PFOS | AB |
| Figure S6Q | Welch's ANOVA with post-hoc Games-Howell's multiple comparisons test | see figure | Control = 130-190; PFOS = 88-99 | 5 dpf | 0.1% DMSO; 7 µM PFOS; 14 µM PFOS | AB |
| Figure S6R | Welch's ANOVA with post-hoc Games-Howell's multiple comparisons test | see figure | Control = 130-190; PFOS = 88-99 | 5 dpf | 0.1% DMSO; 7 µM PFOS; 14 µM PFOS | AB |
| Figure S7A | Two-Way ANOVA with post-hoc Sidak multiple comparisons test | Light off = 0.0139 | Control = 92; PFOS = 97 | 3 dpf | 0.1% DMSO; 7 µM PFOS | AB |
| Figure Suppl 7B | Two-Way ANOVA with post-hoc Sidak multiple comparisons test | ns | Control = 92; PFOS = 97 | 3 dpf | 0.1% DMSO; 7 µM PFOS | AB |
| Figure Suppl 7C | Two-Way ANOVA with post-hoc Sidak multiple comparisons test | Light on = 0.0075; Light off = <0.0001 | Control = 100; PFOS = 100 | 3 dpf | 0.1% DMSO; 14 µM PFOS | AB |
| Figure Suppl 7D | Two-Way ANOVA with post-hoc Sidak multiple comparisons test | Light on = 0.0015; Light off = 0.0018 | Control = 100; PFOS = 100 | 3 dpf | 0.1% DMSO; 14 µM PFOS | AB |
| Figure Suppl 7E | Two-Way ANOVA with post-hoc Sidak multiple comparisons test | Light off = 0.0061 | Control = 88; PFOS = 84 | 3 dpf | 0.1% DMSO; 28 µM PFOS | AB |
| Figure Suppl 7F | Two-Way ANOVA with post-hoc Sidak multiple comparisons test | ns | Control = 88 PFOS = 84 | 3 dpf | 0.1% DMSO; 28 µM PFOS | AB |
| Figure Suppl 7G | Two-Way ANOVA with post-hoc Sidak multiple comparisons test | Light on = 0.0004 | Control = 102; PFOS = 98 | 5 dpf | 0.1% DMSO; 7 µM PFOS | AB |
| Figure Suppl 7H | Two-Way ANOVA with post-hoc Sidak multiple comparisons test | Light on = <0.0001 | Control = 102; PFOS = 98 | 5 dpf | 0.1% DMSO; 7 µM PFOS | AB |
| Figure Suppl 7I | Two-Way ANOVA with post-hoc Sidak multiple comparisons test | Light on = <0.0001; Light off = <0.0001 | Control = 97; PFOS = 89 | 5 dpf | 0.1% DMSO; 14 µM PFOS | AB |
| Figure Suppl 7J | Two-Way ANOVA with post-hoc Sidak multiple comparisons test | Light on = <0.0001; Light off = 0.0004 | Control = 97; PFOS = 89 | 5 dpf | 0.1% DMSO; 14 µM PFOS | AB |
| Figure S8B | Non-parametric ANOVA (Kruskal-Wallis test) with Dunn's test for multiple comparisons | see figure | 20-22 per group | 3 dpf | 0.1% DMSO; 7 µM PFOS; 14 µM PFOS; 28 µM PFOS | AB |
| Figure S8C | Non-parametric ANOVA (Kruskal-Wallis test) with Dunn's test for multiple comparisons | see figure | 20-22 per group | 5 dpf | 0.1% DMSO; 7 µM PFOS; 14 µM PFOS | AB |
| Figure 5B | Unpaired parametric t-test with Welch's Correction | 0.0237 | 14-22 per group | 5 dpf | 0.1% DMSO; 7 µM PFOS | *Tg*(*elavl3:CaMPARI)* |
| Figure 5C | Two-Way ANOVA with post-hoc Tukey multiple comparisons test | see figure | 13-16 per group | 5 dpf | 0.1% DMSO; 7 µM PFOS | *Tg*(*elavl3:CaMPARI)*  *irf8* WT and *irf8* Mutant |
| Figure 5D | Two-Way ANOVA with post-hoc Tukey multiple comparisons test | see figure | 13-16 per group | 5 dpf | 0.1% DMSO; 7 µM PFOS | *Tg*(*elavl3:CaMPARI)*  *irf8* WT and *irf8* Mutant |
| Figure 5E | Two-Way ANOVA with post-hoc Tukey multiple comparisons test | see figure | 13-16 per group | 5 dpf | 0.1% DMSO; 7 µM PFOS | *Tg*(*elavl3:CaMPARI)*  *irf8* WT and *irf8* Mutant |
| Figure 5F | Two-Way ANOVA with post-hoc Tukey multiple comparisons test | see figure | 13-16 per group | 5 dpf | 0.1% DMSO; 7 µM PFOS | *Tg*(*elavl3:CaMPARI)*  *irf8* WT and *irf8* Mutant |
| Figure 5G | Two-Way ANOVA with post-hoc Tukey multiple comparisons test | see figure | 13-16 per group | 5 dpf | 0.1% DMSO; 7 µM PFOS | *Tg*(*elavl3:CaMPARI)*  *irf8* WT and *irf8* Mutant |
| Figure S9A | Two-Way ANOVA with post-hoc Tukey multiple comparisons test | See figure | 14-22 per group | 3 dpf | 0.1% DMSO; 28 µM PFOS | AB  *irf8* WT and *irf8* Mutant |
| Figure S9B | Two-Way ANOVA with post-hoc Tukey multiple comparisons test | See figure | 14-22 per group | 3 dpf | 0.1% DMSO; 28 µM PFOS | AB  *irf8* WT and *irf8* Mutant |
| Figure S9C | Two-Way ANOVA with post-hoc Tukey multiple comparisons test | See figure | 14-22 per group | 3 dpf | 0.1% DMSO; 28 µM PFOS | AB  *irf8* WT and *irf8* Mutant |
| Figure S9D | Two-Way ANOVA with post-hoc Tukey multiple comparisons test | See figure | 14-22 per group | 3 dpf | 0.1% DMSO; 28 µM PFOS | AB  *irf8* WT and *irf8* Mutant |
| Figure S9E | Two-Way ANOVA with post-hoc Tukey multiple comparisons test | See figure | 14-22 per group | 5 dpf | 0.1% DMSO; 7 µM PFOS | AB  *irf8* WT and *irf8* Mutant |
| Figure S9F | Two-Way ANOVA with post-hoc Tukey multiple comparisons test | See figure | 14-22 per group | 5 dpf | 0.1% DMSO; 7 µM PFOS | AB  *irf8* WT and *irf8* Mutant |
| Figure S9G | Two-Way ANOVA with post-hoc Tukey multiple comparisons test | See figure | 14-22 per group | 5 dpf | 0.1% DMSO; 7 µM PFOS | AB  *irf8* WT and *irf8* Mutant |
| Figure S9H | Two-Way ANOVA with post-hoc Tukey multiple comparisons test | See figure | 14-22 per group | 5 dpf | 0.1% DMSO; 7 µM PFOS | AB  *irf8* WT and *irf8* Mutant |
| Figure 6I | Two-Way ANOVA with post-hoc Sidak multiple comparisons test | see figure | 14-20 per group | 3 dpf | 0.1% DMSO; 28 µM PFOS | *Tg(elavl3:Gal4;cryaa:RFP;*  *UAS:eNpHR3.0;mpeg1:EGFP)* |
| Figure S10D | Non-parametric ANOVA (Kruskal-Wallis test) with Dunn's test for multiple comparisons | see figure | 14-21 per group | 3 dpf | 0.1% DMSO; 28 µM PFOS; 5 mM PTZ | Tg(*mpeg1:EGFP*) |
| Figure 7C | Non-parametric ANOVA (Kruskal-Wallis test) with Dunn's test for multiple comparisons | see figure | 21 per group | 5 dpf | 0.1% DMSO; 64 µM PFOA | Tg(*elavl3:CaMPARI*) |
| Figure 7F | Unpaired non-parametric t-test (Mann-Whitney Wilcoxon) | 0.2359 | 8-9 per group | 3 dpf | 0.1% DMSO; 64 µM PFOA | Tg*(mpeg1:EGFP)* |
| Figure Suppl 11B | Unpaired parametric t-test with Welch's Correction | 0.1528 | Control = 24; PFOA = 46 | 5 dpf | 0.1% DMSO; 64 µM PFOA | Tg(*elavl3:CaMPARI*) |
| Figure Suppl 11C | Unpaired parametric t-test with Welch's Correction | 0.6189 | Control = 24; PFOA = 46 | 5 dpf | 0.1% DMSO; 64 µM PFOA | Tg(*elavl3:CaMPARI*) |
| Figure Suppl 11D | Unpaired parametric t-test with Welch's Correction | 0.166 | Control = 24; PFOA = 46 | 5 dpf | 0.1% DMSO; 64 µM PFOA | Tg(*elavl3:CaMPARI*) |
| Figure Suppl 11E | Two-Way ANOVA with post-hoc Sidak multiple comparisons test | See figure | Control = 37; PFOA = 35 | 5 dpf | 0.1% DMSO; 64 µM PFOA | Tg(*elavl3:CaMPARI*) |
| Figure Suppl 11F | Two-Way ANOVA with post-hoc Sidak multiple comparisons test | See figure | Control = 37; PFOA = 35 | 5 dpf | 0.1% DMSO; 64 µM PFOA | Tg(*elavl3:CaMPARI*) |
| Figure Suppl 11G | Unpaired parametric t-test with Welch's Correction | ns | Control = 37; PFOA = 35 | 5 dpf | 0.1% DMSO; 64 µM PFOA | Tg(*elavl3:CaMPARI*) |
| Figure S12C | Unpaired parametric t-test with Welch's Correction | <0.0001 | 5-8 per group | 5 dpf | 0.1% DMSO; 64 µM PFOA | Tg(*elavl3:CaMPARI*) |
